# Explainable machine learning identifies features and thresholds predictive of immunotherapy response

**DOI:** 10.1101/2025.03.23.643560

**Authors:** Khoa A. Tran, Venkateswar Addala, Lambros T. Koufariotis, Jia Zhang, Scott Wood, Conrad Leonard, Lotte L. Hoeijmakers, Christian U. Blank, Mireia Crispin-Ortuzar, Elizabeth D. Williams, Olga Kondrashova, John V. Pearson, Nicola Waddell

**Affiliations:** Cancer Program, QIMR Berghofer Medical Research Institute; Brisbane, Australia; School of Biomedical Sciences, Queensland University of Technology (QUT); Brisbane, Australia; Department of Medical Oncology, Netherlands Cancer Institute; Amsterdam, The Netherlands; Department of Medical Oncology, Leiden University Medical Center; Leiden, The Netherlands; Department of Oncology, University of Cambridge; Cambridge, UK; Cancer Research UK Cambridge Centre, University of Cambridge; Cambridge, UK; Australian Prostate Cancer Research Centre – Queensland (APCRC-Q) and Queensland Bladder Cancer Initiative (QBCI); Brisbane, 4000, Australia; Centre for Genomics and Personalised Health, Queensland University of Technology (QUT); Brisbane, QLD, Australia

## Abstract

**Background:** Immunotherapy has improved patient survival for multiple cancer types, including melanoma. While a variety of molecular features have been linked to response to immune checkpoint inhibitors (ICI) treatment, clinically established biomarkers, such as tumour mutation burden (TMB) and PD-L1 expression, have shown limitations in accurately categorising responders versus non-responders. Due to the complex nature of ICI response, which includes cancer intrinsic and extrinsic features within the tumour microenvironment (TME), using a single biomarker to predict response is insufficient, necessitating the need to identify accurate clinical and multi-omic molecular predictors.

**Methods:** We integrate clinical, DNA and RNA sequencing data from four datasets, comprising 138 melanoma patients treated with ICI, to develop machine learning models for predicting ICI response. The performance of each model was evaluated using an independent dataset of patients with cutaneous melanoma (n=53). Interactions between trained models and features were rationalised using the explainability method SHAP.

**Results:** The most optimal model was the multi-omic random forest model, with AUC-ROC of 0.78 when predicting response in the independent test dataset. Using SHAP, we predicted thresholds for mutational signatures, neoantigen load, immune cell-type abundance and immune receptor LAG3 expression. The relationship between these influential features and their SHAP scores revealed numerical thresholds constituting good and poor patient response.

**Conclusions:** This approach highlights patient response to ICI is influenced by both cancer intrinsic and extrinsic features, as well as identifies candidate biomarkers that could inform the use of ICI and potentially assist in the selection of combination therapy in melanoma treatment.

## BACKGROUND

The advent of immune checkpoint inhibitors (ICI) has revolutionised treatment of melanoma [1–3] with ICI that block T cell checkpoint molecules, such as programmed cell death protein 1 (PD-1) and cytotoxic T lymphocyte–associated protein 4 (CTLA-4), approved for melanoma treatment. Unfortunately, not all patients respond to ICI [4], as some show varied outcomes at both early and late stages of cancer. A variety of molecular features have been linked to response to ICI treatment. Even so, the clinically established biomarkers, tumour mutation burden (TMB) [5] and expression of PD-L1 [6], have not effectively fulfilled the role of accurately categorising responders versus non-responders, showing only a modest correlation with response to ICI in 22 tumour types [7]. Other markers including neoantigen load [8–11], cytolytic activity [12], histopathology data [13] and gene expression profiles (GEP) [14] have been used to predict effectiveness of ICI therapies. However, due to the complex nature of ICI response, which includes cancer intrinsic and extrinsic features within the tumour microenvironment (TME) [15], using a single biomarker to predict response is not sufficient.

Several studies have utilised DNA and/or RNA sequencing (RNA-Seq) data to predict responses to ICI [8, 10, 11, 16–18]. Subsequently, integration of DNA and RNA-seq data has occurred, including a small integrative study comprising multiple melanoma subtypes to predict ICI response [9], and a pan-cancer analysis that incorporated individual biomarkers into a multivariable predictor that was reported to outperform TMB alone [8]. Nonetheless, these integrated investigations have not been validated in independent samples, or employed limited features that do not adequately represent the intricacies of the tumour and its microenvironment in melanoma.

Here, we integrated five multi-omic datasets that measure tumour intrinsic and extrinsic features from pre-ICI treated melanoma patients to develop an ensemble machine learning approach to predict responses to ICI in melanoma. We validated the accuracy of the predictive models in an independent dataset. The overall machine learning approach is widely applicable to other ICI treated tumour types and can be customised to incorporate tumour-defined features that associate with ICI responses.

## METHODS

### Data used in this study

The DNA and RNA sequence data from 5 published studies [10, 16–19] was accessed in FASTQ format and re-analysed to ensure data uniformity. Four of these datasets [16–19] were comprised of exome and RNA-Seq, while the Newell et al. dataset [10] was whole genome and RNA-Seq. Only patients with matched DNA and RNA sequence data (**Table S1**), age at diagnosis and ICI responses defined in their respective studies were included in this study. To harmonise the patient response across studies, we grouped patients with complete or partial responses as good responders and those with progressive disease as poor responders. Patients classified as stable disease were excluded from the training data and used as test only. Overall, in the current study we assembled 53 patients from Campbell et al. [16] (19 good responders, 26 poor responders, 8 stable disease); 23 patients from Hugo et al. [19] (13 good responders, 10 poor responders); 53 patients (34 good responders, 15 poor responders, four stable) from Newell et al. [10]; 42 patients from Riaz et al. [17] (10 good responders, 17 poor responders, 15 stable disease) and 58 patients from Rozeman et al. [18] (41 good responders, 17 poor responders) (**Table S1**).

### Machine learning

#### Feature selection

The clinical features were patient age at diagnosis, patient biological sex and binary mutation status of *BRAF*, *KRAS/NRAS* or *NF1* (“Mutated” or “Not mutated”). The DNA features were selected to capture cancer intrinsic features that may be predictive of ICI response [15] and included tumour HLA zygosity, TMB, neoantigen load, tumour purity, tumour ploidy, and mutational signature 7 linked to UV light and sun exposure that have been previously described in melanoma [20, 21]. Features from RNA-Seq data were selected to represent cancer extrinsic properties of the TME that may be indicative of ICI response [15]. These features included the gene expression level of 16 immune checkpoint receptors, four immune gene signatures previously reported to be associated with ICI response[14], two immune scores (TIDE [22] and cytolytic score [12]), as well as the relative proportion of 22 immune cells within the TME. A detailed description of the features and the process to generate these features is included (**Supplementary Note 1**).

#### Model architecture

We used scikit-learn v1.3.2 to build five machine learning classifiers: logistic regression (LR), random forest (RF), support vector machine (SVM), an ensemble comprising of LR and RF (Ensemble:LR+RF), and an ensemble comprising of LR, RF and SVM (Ensemble:LR+RF+SVM). The two ensemble models employed soft voting (**Supplementary Note 2**). Before any data ingested into a machine learning model during both training and test, we also applied one-hot encoding on categorical variables and z-score scaling on numerical features (as described in **Supplementary Note 2**).

#### Model training, hyperparameter optimisation, and test prediction

We trained each machine learning model using five-fold cross-validation to optimise each model’s hyperparameters, with area under the curve-receiver operating characteristic (AUC-ROC) as the objective metric. Additionally, we repeated this cross-validation process ten times with different randomisation in creating the five “folds”, resulting in 50 AUC-ROC scores for each hyperparameter combination. The hyperparameter setting with the highest average AUC-ROC for each model was selected to refit each model on the entire training dataset. For the LR, RF, SVM and Ensemble:LR+RF models, we used the exhaustive *GridSearchCV()* function of scikit-learn and checked all possible settings. For the Ensemble:LR+RF+SVM model, we used the *RandomSearchCV(steps=10,000)* function. A list of searched hyperparameters for each model and their final settings are provided in **Supplementary Note 2**. Refitted models with the best hyperparameters were then used to predict ICI response in test samples. It is important to note that in the experiment where each dataset besides Newell et al. [10], (Campbell et al. [16], Hugo et al. [19], Riaz et al. [17], and Rozeman et al. [18]) was iteratively used as the test data, we only retained cutaneous and non-stable-disease samples for the corresponding training and test dataset.

#### Features selection

We defined features as either clinical, cancer intrinsic (DNA), or cancer extrinsic (RNA). Before either training or test data was ingested into the machine learning pipeline described above, we conducted a Mann-Whitney U-test using training data (null hypothesis: no difference between good versus poor responders, i.e. *p*-value>0.05). Features with *p*-value ≤ 0.05 were selected. Pair-wise correlations among the remaining features were computed using Pearson’s *r* (for pairs of numerical-numerical features), and Eta Correlation Ratio (for pairs of numerical-categorical features). Notably, Sex was the only selected categorical feature, hence there were no pairs of categorical-categorical features.

We performed hierarchical clustering on the features using the dendrogram function in the Python package plotly. To stay consistent with the data transformations applied in the machine learning pipeline, we ensured that the same steps were followed for the hierarchical clustering, i.e., log2p1/log10 for relevant features and z-score scaling on all numerical features (**Supplementary Note 1)**. Categorical features (i.e., sex) were kept as is. By manually annotating the hierarchical clustering output, we identified groups of highly correlated (collinear) features.

We trained the five machine learning models (LR, RF, SVM, Ensemble:LR+RF, and Ensemble:LR+RF+SVM) using all features that passed the significant-association-with-ICI test. For each trained model, we calculated the median absolute SHAP score for each feature using only the correctly predicted samples in the training data. Within each collinear cluster identified using the methodology described above, only one feature with the highest median absolute SHAP score was retained. The remaining features were subsequently used for hyperparameter optimisation, training and test evaluation.

#### Performance evaluation metrics

Where appropriate, we used AUC-ROC and F1 to evaluate machine learning models’ performance. Of note, both AUC-ROC and F1 are sensitive to class imbalance. Therefore, we chose to use the “weighted” version of these metrics, by which they are is defined as

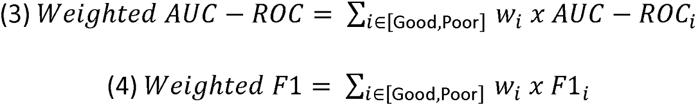

where AUC-ROC*_i_* and F*_i_* are metrics calculated from either good or poor response predictions, and *w_i_* is the corresponding proportion of patients in each class. Of note, this is a binary classification (i.e. good vs poor ICI response), hence Weighted AUC-ROC is analogous to Micro and Macro AUC-ROC.

#### Features contribution in test samples

We implemented SHAP on test samples for the machine learning models trained only with selected features as described in *Features selection*. We used the KernelExplainer [23] to explain the LR, SVM, Ensemble:LR+RF, and Ensemble:LR+RF+SVM models. We used the TreeExplainer [24] to explain the RF model, which is designed for tree-based machine learning models.

## RESULTS

### Multi-omic machine learning workflow

We developed a multi-omic machine learning workflow to predict whether individual melanoma patients would have a good or poor response to ICI. Good response was assigned when patients had a complete or partial response to ICI, poor included patients with progressive disease in response to ICI. Features to train and test the model were extracted from five datasets [10, 16–19] derived from melanoma patients treated with ICI (**Fig. 1A**). The specific ICI treatment was not consistent across patients, reflecting real world clinical care, with n=116 patients receiving single agent ICI (anti-PD1 n=102 or anti-CTLA4 n=14) while n=113 patients received both anti-PD1 and anti-CTLA4 (**Table S1**). We assembled a total of 57 features from each patient that were used to train and test the model (**Supplementary Note 1**). The features consisted of clinical information (patient biological sex, age at diagnosis and *BRAF*, *K/N-RAS*, *NF1* mutation status), tumour tissue DNA sequencing (to capture eight cancer intrinsic features) and RNA-Seq (to capture 44 tumour extrinsic features) (**Fig. 1A**). A subset of features was selected to train and test each predictive mode (**Fig. 1B**). Four of the five datasets [16–19] were used to train the model (**Fig. 1C**). Patients with non-cutaneous melanoma (n=23) or cutaneous samples with stable disease (n=15) were excluded from training and withheld as additional test data. The selected features from each modality (clinical, DNA or RNA) were used to train single- and multi-omic machine learning models comprised of Logistic Regression (LR), Random Forrest (RF) and Support Vector Machines (SVM) as well as ensemble approaches (LR+RF and LR+RF+SVM) (**Fig. 1C**). The performance of each model was assessed and validated using an independent dataset [10] comprising 49 cutaneous melanoma samples. In addition, non-cutaneous melanoma samples (n=15), non-cutaneous melanoma samples with stable disease (n=8) and cutaneous samples with stable disease (n=19) which were excluded from the training data [16, 17] were used to assess the performance of each model (**Fig. 1d**). We used an explainability approach, SHapley Additive exPlanations (SHAP) [23, 24], to identify features associated with ICI response.

**Fig. 1:**
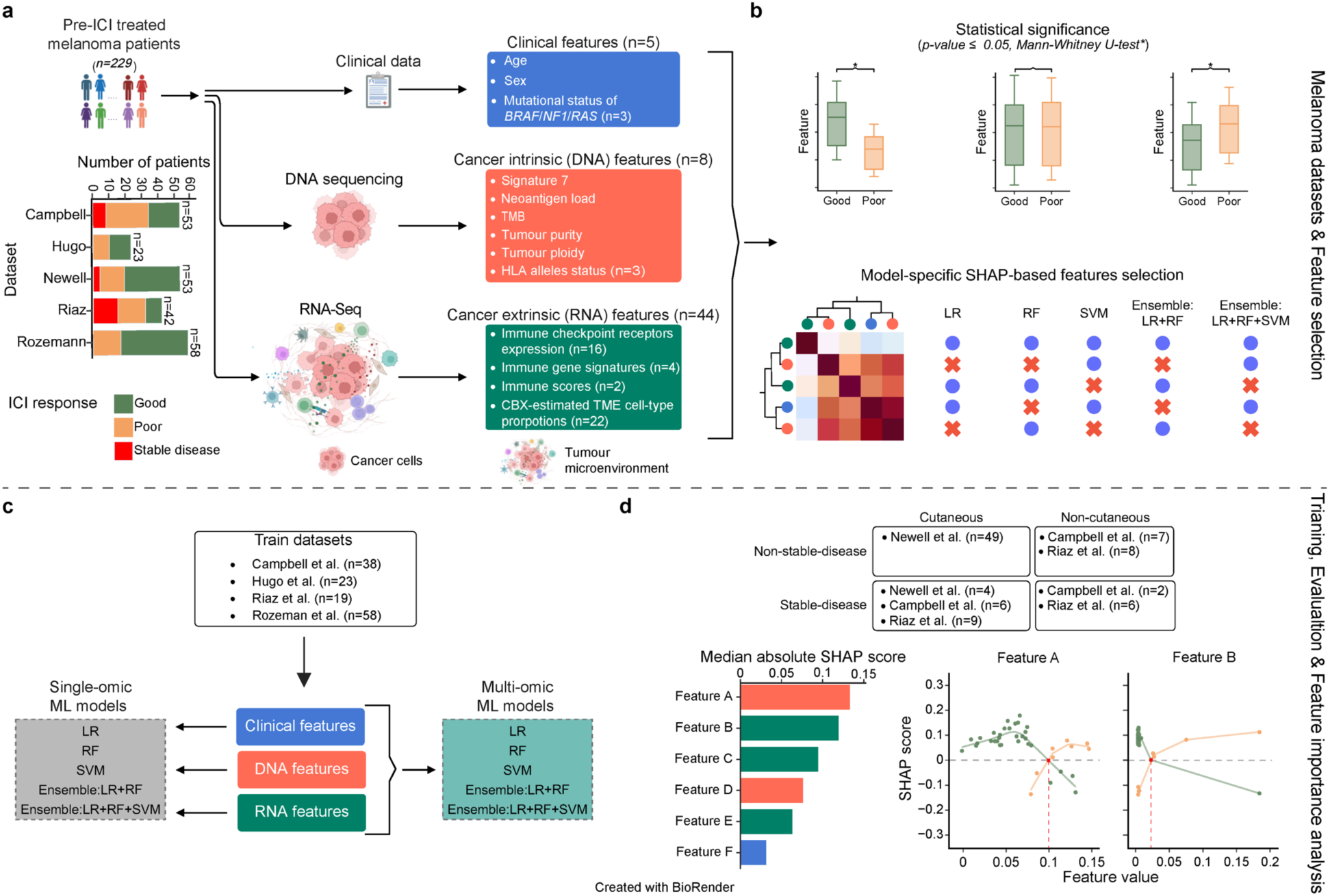
Experimental design overview. **A)** Data from 229 pre-ICI treated melanoma patients stratified into good (green), poor (orange) response to ICI or stable disease (red) were collected from 5 datasets. Each dataset included clinical features (n=5), cancer intrinsic features extracted from DNA sequencing (n=8) and cancer extrinsic features from RNA-Seq (n=44). Clinical features (blue) include sex of patient, age at diagnosis and mutation status of *BRAF*/*NF1/RAS* genes. Cancer intrinsic (DNA) features (orange) include signature 7, neoantigen load, TMB, tumour purity, tumour ploidy and tumour HLA allele status. Cancer extrinsic (RNA) features (green) include gene expression of immune checkpoint receptors (n=16), immune gene signatures (n=4), immune scores (n=2) and the relative proportion of immune cells (n=22) within the TME. **B)** Features significantly associated with good versus poor response within the training dataset were selected and underwent hierarchical clustering and pairwise correlation analysis. A single representative feature from highly correlated features was selected for each model. We created five machine learning models (LR, RF, SVM and two ensemble models LR+RF and LR+RF+SVM). **C)** Data for cutaneous melanomas from 4 datasets (excluding stable disease) was used to train each model. In single-omic machine learning models, clinical, cancer intrinsic (DNA) and extrinsic (RNA) features are trained separately, whereas in multi-omic ensemble machine learning models features are combined. **D)** Trained models were evaluated using an independent cutaneous melanoma dataset from Newell et al. (n=53, including n=49 with good/poor response and n=4 with stable disease) and additional samples with stable disease and/or non-cutaneous melanoma. Feature importance determined by SHAP scores identified key features differentiating good and poor responders. ICI: immune checkpoint inhibitor, TMB: tumour mutation burden, HLA: human leukocyte antigen, LR: logistic regression, RNA-Seq: RNA sequencing, RF: random forest, SVM: support vector machine, ML: machine learning, SHAP: SHapley Additive exPlanations, TME: tumour microenvironment.

### Curating multi-omic features linked to ICI response

We undertook a biology-driven approach to assemble features for the prediction of ICI response. We identified 57 features and extracted data for these within the training (n=138) [16–19] (**Fig. 2** and **Table S1**) and independent test (n=53) [10] (**Fig. S1** and **Table S1**) datasets. To reduce the number of features we undertook multiple consecutive feature selection approaches (**Fig. 1B**). In the first instance, we selected from the 57 features those that were significantly different between patients with a good and poor response within the training data (*P*≤0.05, according to Mann-Whitney U-test (MWU), **Table S2**). Although the clinical features were not associated with ICI response across the training data, in the Rozeman et al. dataset [17], older patients were more likely to respond to ICI (*P*=0.05, MWU), while not significant this trend was stronger for male patients compared to female patients (**Fig. S2A**). Previous reports had linked age [25] and sex [26] to ICI response, therefore we elected to retain these clinical features. Of the eight DNA features, only the proportion of mutational signature 7 was significantly associated (*P=*0.04, MWU) with ICI response (**Fig. S2B** and **Table S2**). In addition, although not significantly associated with ICI response in the training data, we included TMB and neoantigen load in the predictive model. TMB is an FDA approved marker for ICI treatment [5] and was significantly higher in good responders within the Rozeman et al. dataset [18] (*P*=0.02, MWU, **Fig. S2B**), while there are multiple studies supporting the predictive value of neoantigen load [8–11]. Seventeen of the RNA features were significantly associated with ICI response in the training data. These included expression of six genes encoding immune checkpoint receptors (**Fig. S3A**), the cytolytic immune score [12] (*P*=0.04, MWU) and four immune gene signatures (Effector T cell *P*=0.03, and IFNg-6/Effector T cell *P*=0.04, IFNg-6 *P*=0.03 and IFNg expanded 18 *P*=0.01; MWU) (**Fig. S3B**), and the relative proportion of six immune cell types (naïve B cells *P*=3×10^-4^, M1 macrophages *P*=0.04, M2 macrophages *P*=3×10^-3^, CD8 T cells *P*=9×10^-3^, follicular helper T (Tfh) cells *P*=10^-3^, regulatory T (Treg) cells *P*=0.03; MWU) (**Fig. S3C**).

**Fig. 2:**
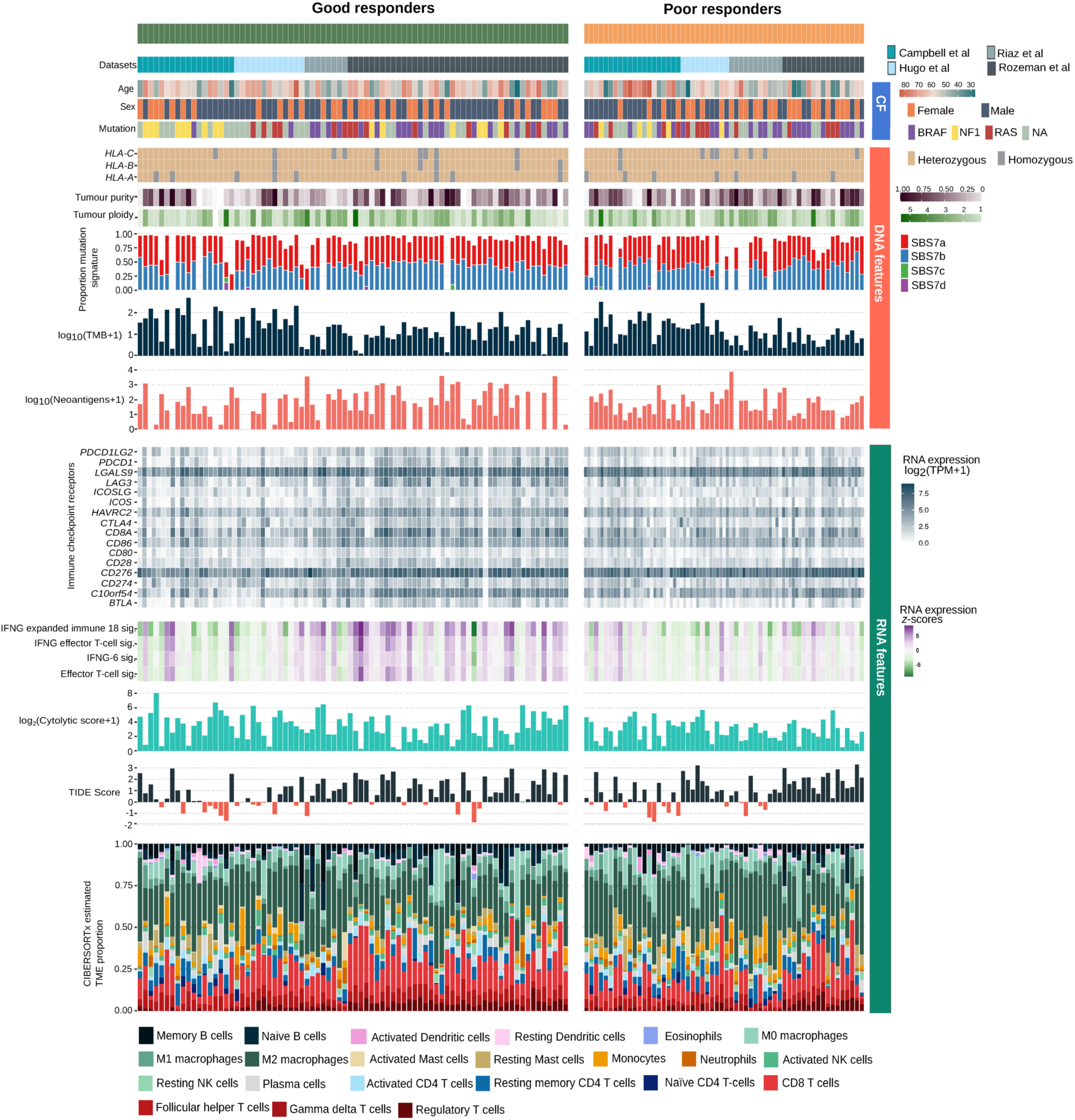
Clinical, DNA and RNA features within the train data. Overview of features from 138 patients within the training data sorted by patient response to ICI, with good (green) and poor (orange) responses. Samples are subsequently sorted by dataset (Campbell et al. Hugo et al., Riaz et al. and Rozeman et al.). The clinical features are: patient age at diagnosis (in years), sex (female, male) and mutation status of *BRAF*, *K/N-RAS* and *NF1* (NA indicates samples that have no identifiable mutation in these genes). Cancer intrinsic (DNA) features are: zygosity of tumour HLA allele, tumour purity (proportion of cancer cells), tumour ploidy, proportion of ultra-violet light mutation signatures (SBS7a, SBS7b, SBS7c, SBS7d), log10p1 transformed neoantigen load and log10p1 transformed TMB. Cancer extrinsic (RNA) features are: log2p1 transformed TPM expression of selected immune checkpoint receptors, gene set signatures scores (Effector T-cell, IFNg-6, a combined IFNg/effector T cell, and an 18-gene expanded IFNg signatures), immune scores (log2p1 transformed cytolytic score and TIDE score) and the proportion of 22 immune cells estimated by deconvolution of RNA-seq data by CIBERSORTx. ICI: immune checkpoint inhibitor, CF: clinical features, HLA: human leukocyte antigen, IFNg: interferon gamma, log2p1: logarithm base 2 of value plus 1, log10p1: logarithm base 10 of value plus 1, NK: natural killer, TMB: tumour mutation burden, TIDE: T-cell immune dysfunction and exclusion.

The 22 selected features were used to train five supervised machine learning models (**Fig. 1C**). We used SHAP scores to quantify the influence of each feature to accurately predict ICI response. To account for collinear features, we kept a representative feature from four collinear clusters (**Fig. 3A**) and retrained each model using features with the highest median absolute SHAP scores (**Fig. 3B**). This resulted in 15 features selected for each model, comprising two clinical features (age and sex) plus a combination of two DNA features (signature 7, neoantigen load or TMB) and 11 RNA features comprising the expression of three or four marker genes (*CD28*, plus *BTLA4* or *ICOS*, plus *PDCD1* or *LAG3*, plus *CD8A* or IFNg expanded 18), one or two immune score (IFNg-6, plus IFNg expanded 18 or *CD8A*) and six cell type proportions.

**Fig. 3:**
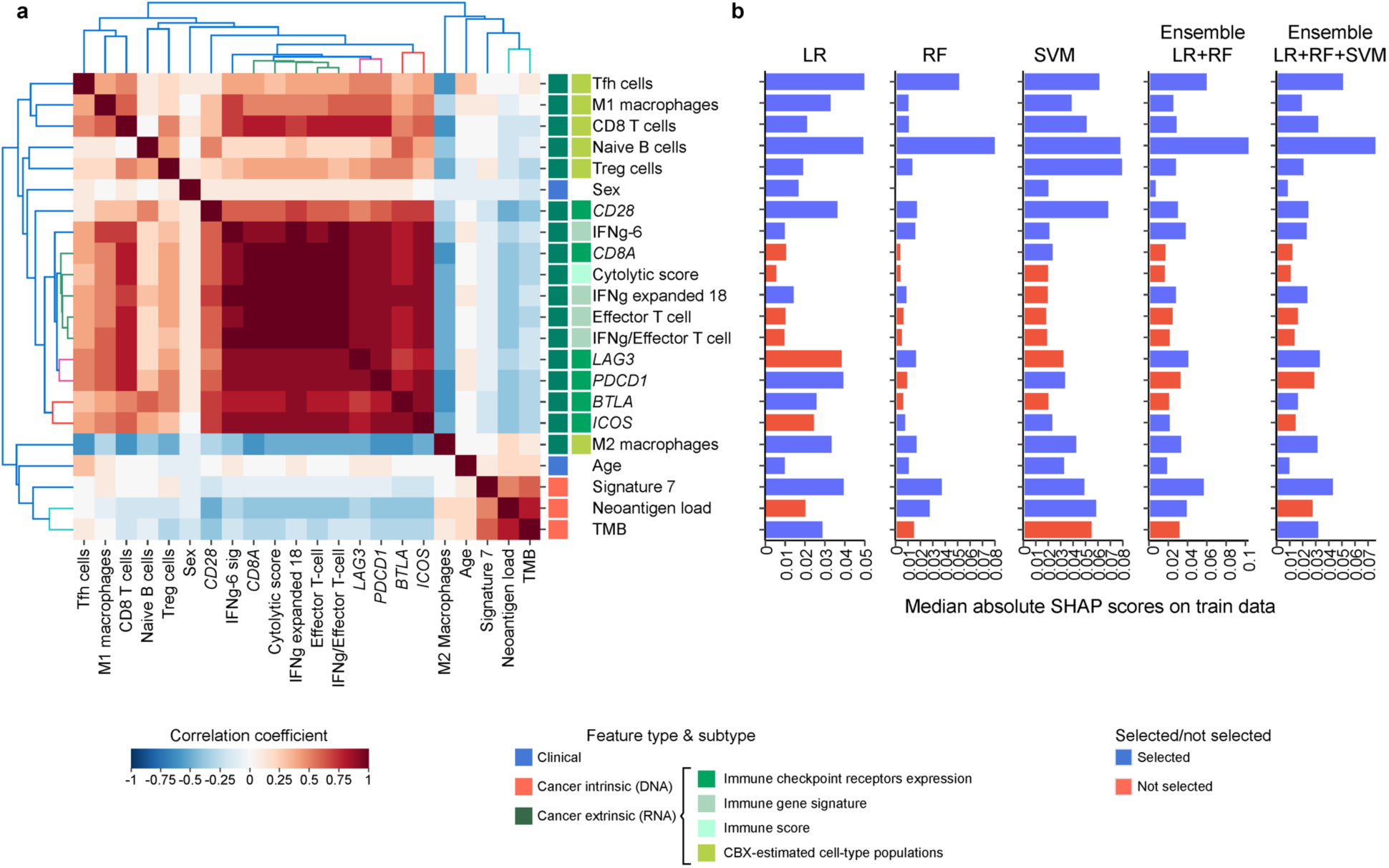
Identification of features multi-collinearity and model-specific feature selection. **A)** Correlation plot showing unsupervised clustering of the correlation between each feature. Numerical-numerical pairwise correlation for each feature pair is based on Pearson’s r coefficient. Categorical-numerical pairwise correlation, e.g. between neoantigen load and biological sex, was based on Eta Correlation Ratio. Scale indicates the correlation, with darker shade of blue representing stronger negative correlation, and darker shade of red representing stronger positive correlation. Four sets of highly correlated features are indicated by coloured clusters (green, pink, red, and light blue cluster). Each feature is colour coded by the type of feature (Clinical: blue, cancer intrinsic (DNA): orange and cancer extrinsic (RNA): green). **B)** Median absolute SHAP scores (x-axis) computed over training samples for each feature (y-axis) after training each model on the feature set. For each model, blue bars represent selected features, and red bars show features discarded among the four highly correlated feature clusters. IFNg: interferon gamma, LR: logistic regression, RF: random forest, SVM: support vector machine, SHAP: Shapley Additive exPlanations, Tfh cells: follicular helper T cells, TMB: tumour mutation burden, Treg cells: regulatory T cells.

### Multi-omic machine learning approach is best predictor of ICI response

Each of the five machine learning models trained on their respective 15 SHAP-selected features were cross-validated within the training data (n=138) (**Fig. 4A**). The RF model exhibited the most optimal training performance for predicting ICI response, evidenced by the highest median values and the narrowest interquartile ranges for F1 and AUC-ROC across most single- and multi-omic feature sets (**Fig. 4B**). When the model was trained with single-omic feature sets the median AUC-ROC score for training data was 0.44 for clinical features, 0.69 for DNA features and 0.72 for RNA features. Performance improved when the multi-omic data was integrated, with AUC-ROC scores of 0.70 for clinical+DNA features, 0.69 for clinical+RNA features, 0.74 for DNA+RNA features, and 0.72 for clinical+DNA+RNA features (**Fig. 4B**).

**Fig. 4:**
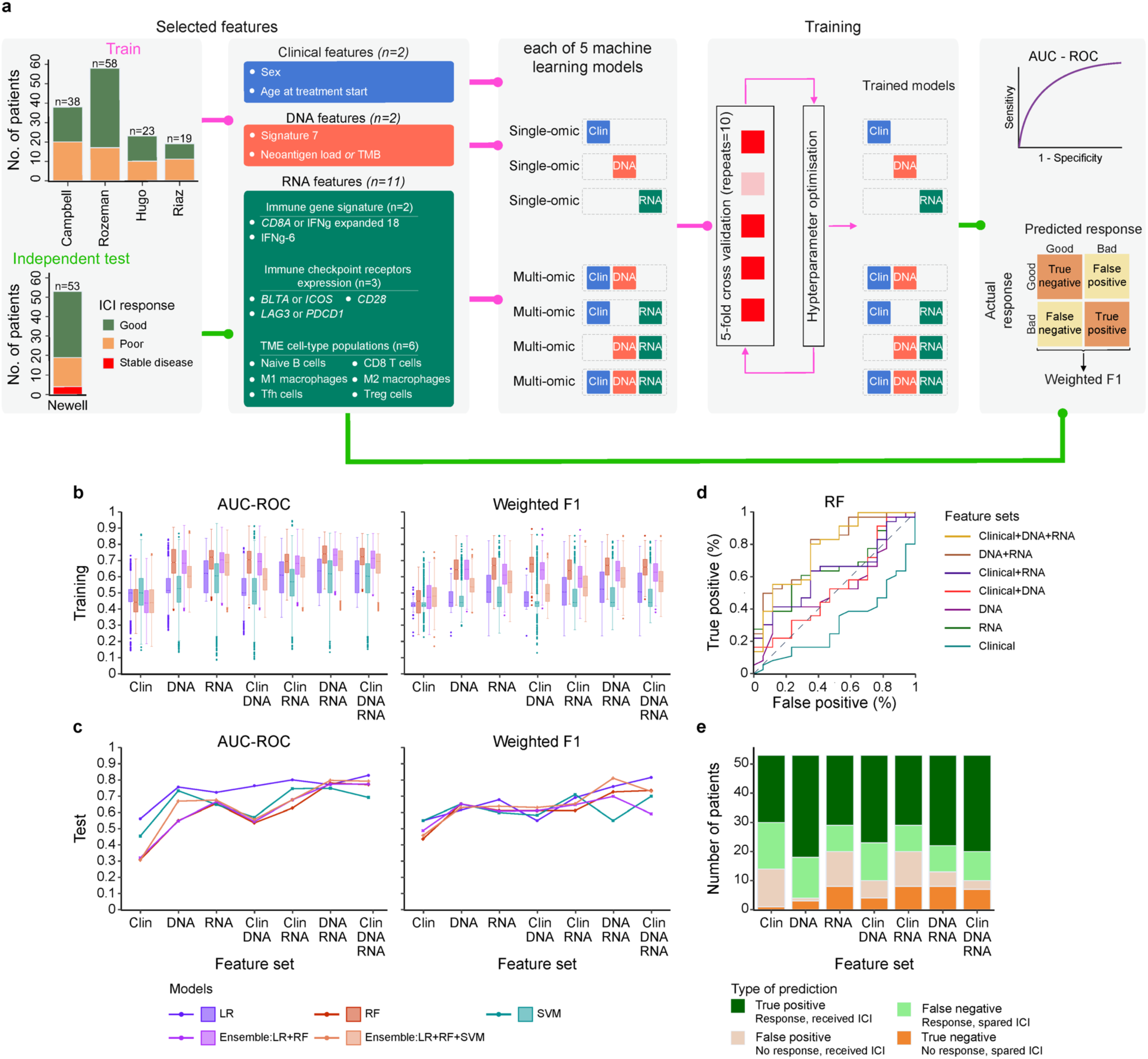
Single- and multi-omic machine learning predictor of immune checkpoint inhibitor response. **A)** Schematic of machine learning framework with training data comprising 138 patients with cutaneous melanoma from four datasets and an independent test dataset of patients with cutaneous melanoma (n=53). Clinical features (blue, n=2), cancer intrinsic features from DNA (orange, n=2) and cancer extrinsic features from RNA (green, n=11) used by the machine learning models are shown. The features were used in the model as single-(n=3) or multi-omic (n=4) sets. Model predictions on both training and test data were assessed using AUC-ROC and weighted F1 scores. **B)** Machine learning models’ performance during training. Boxplots depict AUC-ROC and weighted F1 distributions across five-fold cross-validation, repeated ten times. **c)** Machine learning models’ performance during testing. Line plots depict single AUC-ROC and weighted F1 scores of the best models, based on training performance, on the test data. For all boxplots **(B)** and line plots **(C)**, higher AUC-ROC/weighted F1 values indicate better performance. **D)** ROC curves of the RF model trained on different single and multi-omic feature sets achieved from the independent test cohort. The dashed line at 45-degree angle indicates random prediction. **E)** Stacked bar plots showing the number of patients (from the total n=53) predicted to have a good response (true positive) or a poor response (true negative) to ICI, as well as the number of patients that would receive ICI with no response (false positive) or would not receive ICI but might have benefited (false negative). AUC-ROC: area under the curve of the receiver operating characteristic curve, Clin: Clinical, ICI: immune checkpoint inhibitor, LR: logistic regression, RF: random forest, SVM: support vector machine.

The top performing RF model was then evaluated using an independent test dataset [10] comprising 53 cutaneous samples, which includes 49 samples categorised as good or poor response and 4 samples as stable disease (**Fig. 4A**). The integrated multi-omic features had the best performance with an AUC-ROC score of 0.78 for DNA+RNA features and weighted F1 score of 0.73 (**Fig. 4C**). Within the test data, the number of true positive (patient correctly predicted to have a good response to ICI) and true negative predictions (patient correctly predicted to have a poor response to ICI) was higher when the models utilised the multi-omic features (**Fig. 4D** and **E**). Using clinical features alone would have resulted in 24.53% (13 out of 53) of patients receiving ICI needlessly as they would not respond (false positive) and 30.19% (16 out of 53) of patients not given ICI when they might have benefited from treatment (false negative). In contrast, using the multi-omic features (DNA and RNA) resulted in 73.58% (39 out of 53) accurately receiving ICI or being spared ICI therapy (**Fig. 4E**). This highlights patient response to ICI is multifaceted and a holistic approach to profile tumour samples is needed.

To determine if the training/test data selection impacted ICI response prediction, we repeated the entire feature selection and machine learning workflow using cutaneous and non-stable-disease samples in each dataset as the independent test data, while the remaining four data sets served as training data. This resulted in SHAP-selected features for each train-test permutation and model (**Fig. S4)**. Similar to using Newell et al. as test data, RF models and multi-omic (DNA+RNA) models consistently achieved the best training performance (**Fig. S5**). However, the test results varied, with AUC-ROC for the multi-omic (DNA+RNA) RF model ranging between [0.51-0.74]. The most promising test results was observed when Newell et al. served as test data, particularly for the multi-omic model, which demonstrated higher AUC-ROC scores (0.78, **Fig. 4C-D**) compared to best performance in other test set configurations (0.61 for Campbell et al. (DNA-RF, **Fig. S5A**), 0.76 for Hugo et al. (Clinical+DNA-SVM, **Fig. S5B**), 0.70 for Riaz et al. (Clinical+DNA+RNA-LR and DNA+RNA-LR, **Fig. S5C**), and 0.77 for Rozeman et al. (DNA+RNA-SVM, **Fig. S5D**)). Interestingly, the ratio between good versus poor responders for train data was most balanced when either Newell et al. (80 good, 58 poor) or Rozeman et al. (73 good, 56 poor), suggesting that generalisability is influenced by class imbalance in the train data.

### Predicting response in stable disease and non-cutaneous patients

In the training data, patients with stable disease were excluded from the training data. The independent cutaneous test data from Newell et al. [10] (n=53) included four with stable disease (**Fig. 1d**), these were previously classified as good or poor response by the Newell et al. study [10] using survival time (good n=2, survival of >6 months or poor n=2, survival of <6 months). Encouragingly, the multi-omic ensemble model correctly predicted 17 of 20 patients with a partial response to ICI as good (**Fig. S6A**) and was able to assign the correct good or poor label to three stable disease patients as categorised by Newell et al. **Fig. S6A**). Interestingly, the predicted probabilities for the two stable disease patients categorised as poor responders in Newell et al. was low (**Fig. S6B**). We attribute this to an absence of stable disease patients in the training data, since stable disease patients were deliberately excluded from the training data as these patients represent a clinical classification challenge. This is evidenced by Campbell et al. [16] classifying patients with stable disease as a good response, Riaz et al. [17] as a poor response, while the Newell et al. study [10] used survival time to classify stable disease as good or poor. In further support of this, additional validation results showed that the model predicted three out of six stable disease patients in Campbell et al. [16] as poor responders, and three out of nine patients in Riaz et al. [17] as good responders (**Fig. S6C**). Future molecular profiling of patients with stable disease may result in a better stratification of patient response in this grey area.

To test whether the model is useful in non-cutaneous disease, we used 23 non-cutaneous samples from Campbell et al. [16] and Riaz et al. [17] as an additional test dataset, eight of the patients had stable disease, three a good response to ICI and 12 a poor response. For the eight non-cutaneous patients with stable disease, when using multi-omic features (Clinical+DNA+RNA) six patients were predicted to have a good response, and two a poor response (**Fig. S7A**). Similar to cutaneous cases, the models were not able to consistently predict response for the patients with stable disease. Of the 15 patients with a good or poor response, the RF model using multi-omic features showed better performance than all other models. When using multi-omic features (Clinical+DNA+RNA), the model accurately predicted ICI response for 14 of the 15 non-cutaneous samples with correct predictions for all mucosal, uveal, and samples with an unknown subtype, and only one incorrect prediction for the single acral sample (**Fig. S7B**). This suggests the model may be translatable to other melanoma subtypes, a finding that needs to be confirmed in a larger series of samples.

### Most influential features of patient ICI response

To identify the most relevant features for predicting ICI response, we calculated median absolute SHAP score for each feature in the test data based on predictions produced by the RF model using DNA+RNA features. The five features with the highest SHAP scores were proportions of follicular helper T cell and naïve B cell (RNA), signature 7 and neoantigen load (DNA), and expression of *LAG3* (RNA) (**Fig. 5A**). Notably, these features also exhibited high importance in the training data (**Fig. 3B**). Conversely, proportions of Treg cells, M2 macrophages, and M1 macrophages, the immune signature IFNg expanded 18, and the expression of *ICOS* had the lowest SHAP scores in the test data (**Fig. 5A**).

**Fig. 5:**
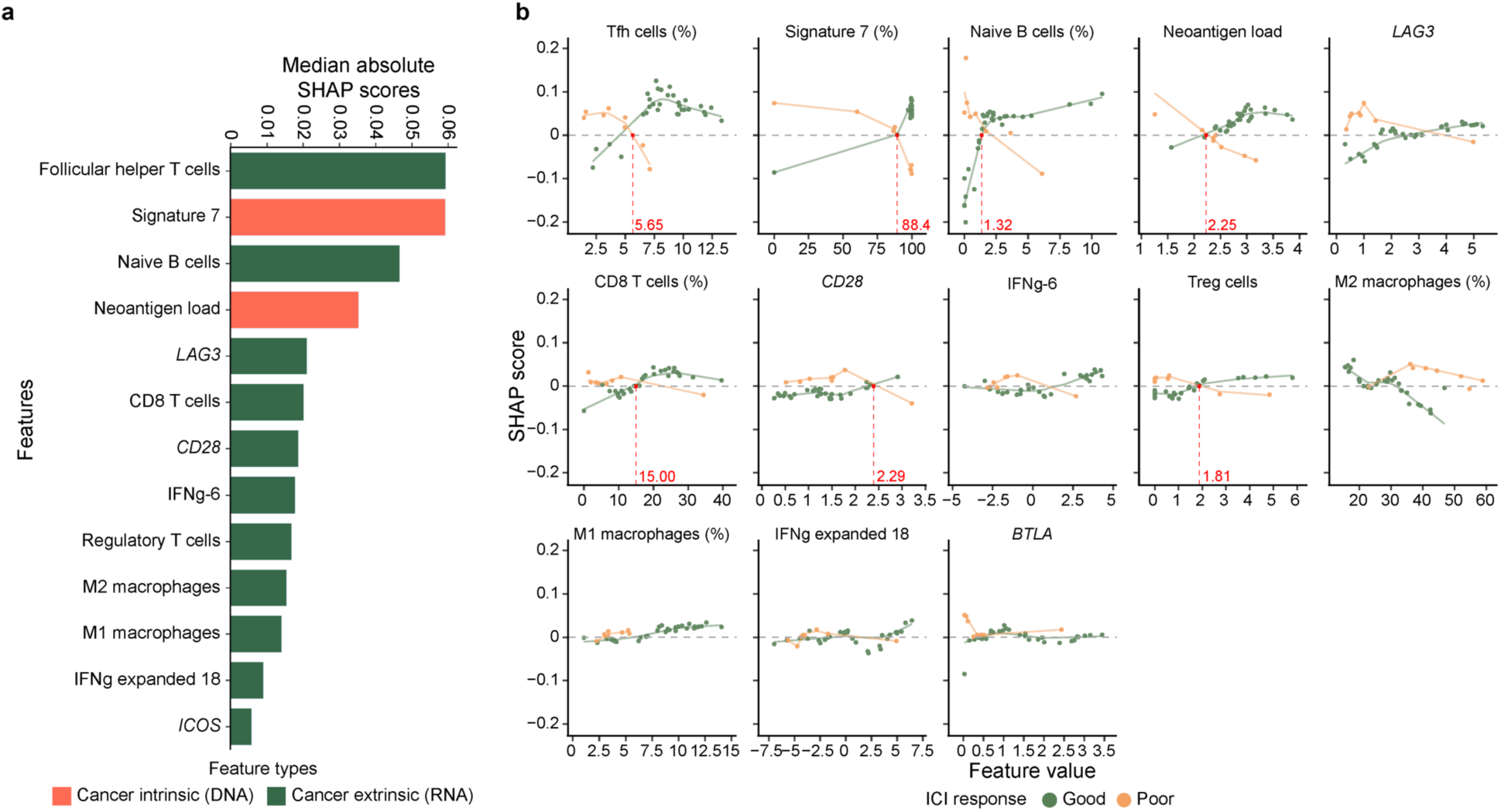
SHAP scores reveal underlying biological drivers of checkpoint inhibitor response. **A)** Median absolute SHAP scores computed over correctly predicted samples in the test data (n=39 out of 53) for each feature based on the RF model trained on multi-omic features (DNA+RNA). The features are coloured by their feature type. The direction of influence (- or + SHAP scores) of each feature is not captured. **B)** Scatter plot of raw SHAP score (y-axis) versus transformed feature values (x-axis) of each feature for patients where the response to ICI was correctly predicted (n=39 out of 53). Neoantigen load is depicted on log10p1 scale, and TPM counts of immune checkpoint receptors (*LAG3*, *CD28*, and *BTLA*) are depicted on log2p1 scale. Each data point represents one patient, coloured by ICI response (good: green and poor: orange). The horizontal dashed line represents the SHAP score of 0, indicating that a feature has no influence on a particular prediction. The green and orange curves represent best fitted curves for, respectively, good and poor responders using locally weighted scatterplot smoothing (LOWESS). Red vertical lines represent identifiable and manually annotated intersections between good (green) and poor (red) LOWESS-fitted curves. ICI: immune checkpoint inhibitor, IFNg: interferon gamma, fSHAP: SHapley Additive exPlanations, Tfh cells: follicular helper T cells, Treg cells: regulatory T cells.

In the samples where ICI response was correctly predicted, we determined whether the SHAP score for each sample was associated with the value of each feature, i.e. would a feature contribute more to response (higher SHAP score) if the feature has a higher value (higher count or expression). Several features showed an association with SHAP score (**Fig. 5B**). Some features contained an inverse relationship of SHAP score and feature value between good and poor responders. For example, good responders with Signature 7 proportions above 88.4% had positive SHAP scores, indicating a supporting influence on response, while those with a Signature 7 proportion below 88.4% had negative SHAP scores, indicating an adversarial influence on response. Conversely, poor responders exhibit an inverse relationship, with positive SHAP scores for Signature 7 proportions below 88.4% and negative SHAP scores for proportions above 88.4%. These findings are suggestive of a distinct feature value threshold that could be used to predict whether a patient is likely to show a good or poor response. Similar relationships were observed in other features; notable examples were neoantigen load (2.25 in log_10_ scale), proportion of Tfh cells (5.65%), proportion of naïve B cells (1.32%), proportion of CD8 T cells (15%), proportion of Treg cells (1.81%) and expression level of *CD28* (2.29 in log_2_ scale) (**Fig. 5B**). This suggests these features have a supporting influence for a good response, but an adversarial influence for poor response. However, it is important to emphasise we interpret these SHAP values as feature influence on a predicted class, not as a singular predictor of ICI response.

As the Clinical+DNA+RNA-LR models achieved the highest F1 scores, i.e., more correct predictions, (**Fig. 4C, Fig. S6**), we utilised SHAP scores of the correctly predicted test samples to explore potential feature thresholds for all test datasets. There were eight significant features selected by SHAP across all permutations: Signature 7, TMB, the proportions of CD8 T cell, Tfh cells, M1 macrophages, and M2 macrophages, and the clinical features age and sex. Interestingly, TMB was consistently among the five features with highest median absolute SHAP scores (**Fig. S8A**). The SHAP-derived feature thresholds of TMB was similar across permutations (1.17-1.43 in log_10_ scale, or 13.79-25.92 in linear scale, **Fig. S8B**). Similar thresholds were observed for the other seven features (56.74-58.17 for age, 81.13-86.08% for Signature 7, 13.19-16.05% for CD8 T cell, 5.70-6.44% for follicular helper T cell, 5.57-6.13% for M1 macrophages, and 29.96-33.63% for M2 macrophage). In terms of sex, SHAP scores indicated that female patients with a good response had a negative SHAP score whereas those with a poor response had a positive SHAP score, and the reverse trend was seen for male patients, except when Rozeman et al. was used as test data (**Fig. S8C**). This finding supports previous literature whereby male melanoma patients respond more favourably to ICI[27]. However, it is important to note that sex was among the least influential feature for both training and test data in three of five train-test permutations (**Fig. 5A**, **Fig. S4, Fig. S8A)**.

## DISCUSSION

Predicting response to ICI is complex and multifaceted, requiring consideration of cancer intrinsic and extrinsic features as well as patient clinical information [15]. We assembled five datasets with clinical, DNA and RNA sequencing data from a diverse group of melanoma patients who received ICI to develop machine learning models to predict ICI response. Using explainable machine learning, we identified the most important features in predicting response to ICI and defined a numerical threshold of response for these features.

The best predictive model was the multi-omic ensemble model combining logistic regression and random forest, with AUC-ROC of 0.78 when predicting response in an independent cutaneous dataset. This model outperforms previous studies that utilised machine learning to predict patient response to ICI using multiple data modalities in melanoma [9] or different cancer types [8]. Liu et al. [9] trained a logistic regression model using a small set of metastatic melanoma patients (n=34 with exome and RNA-Seq) treated with anti-PD-1 (ipilimumab) to predict resistance, achieving an AUC-ROC score of 0.83 in the training data. However, the final predictive model was not validated in an independent dataset. Litchfield et al. [8] used an XGBoost multivariate model trained on 11 biomarkers derived from four cancer types (bladder, head and neck, renal and melanoma) to achieve an AUC-ROC of 0.86 for the prediction of resistance to ICI, however in an independent melanoma dataset [9], it only achieved an AUC-ROC of 0.66, suggesting that the selected markers did not sufficiently capture response to ICI in melanoma.

Accurate prediction relies on the availability of well phenotyped clinical datasets that capture patient variability. A limitation of our study was the number of features and patients available. The data contained few clinical features, and treatment was not standard with some patients receiving different mono or combination ICI treatments. The treatment may impact the model, as in melanoma patients treated with a combination of anti-CTLA-4 and anti-PD-1 the 5-year overall survival is 52%, compared to 44% with anti-PD-1 or 26% with anti-CTLA-4 monotherapy [28]. In terms of cancer extrinsic features, our analysis was limited to specific immune populations and excluded cancer associated fibroblasts and other stromal cell populations which play a role in immunotherapy response [29, 30]. Deconvolution methods which rely on gene expression profiles and cell-type annotations from a single-cell reference, such as BayesPrism [31], may improve TME cellular deconvolution. Additionally, future work to integrate other data modalities, such as the tumour proteome or histopathology information may improve prediction of ICI response.

A notable strength of our study is the utilisation of the explainability method SHAP. We initially used SHAP to select a representative feature from features with high collinearity, subsequently SHAP scores were used to identify the most discriminatory biomarkers of ICI response. The proportion of the UV mutational signature (SBS7) was a discriminatory feature in most models, supporting a previous study. The majority of discriminatory features were RNA features representing cancer extrinsic properties of the TME. The presence of follicular helper T cells was consistently amongst the most discriminatory features in all models. Follicular helper T cells (Tfh) play a crucial role in supporting the maturation and functionality of B cells [32]. In mice with triple-negative breast cancers, a high TMB has previously been linked to activation of B cells mediated by Tfh cells [33] and the presence of both B cells and Tfh cells is associated with longer survival and better therapeutic outcomes in melanoma [34]. Other discriminatory features included immune cells derived from CIBERSORTx deconvolution in descending order of importance, naïve B cells, CD8 T cells, regulatory T cells, M1 and M2 macrophages. B cells have been associated with a favourable response to immunotherapy in multiple cancer types[33, 35, 36]. M1 macrophages play a crucial role in combating tumours by producing pro-inflammatory substances such as TNF-α, IL-1β, and iNOS [37] and expressing high levels of antigen-presenting MHC complexes [38]. Conversely, a high proportion of M2 macrophages is positively associated with a poor response, which aligns with the ability of M2 macrophages to promote tumour growth and cause an immune suppressive environment by excluding T cells from the TME [39]. *LAG3* was the most predictive single gene marker, showcasing the ability of explainable machine learning to identify therapeutic targets. *LAG3* is an ICI target, with the anti-LAG-3 drug relatlimab, approved by the FDA in combination with nivolumab for the treatment of unresectable or metastatic melanoma. Patient response to this treatment (48% of patients with a 12-month progression free survival) [40] is comparable to the anti-CTLA4 and anti-PD-1 combination (ipilimumab/nivolumab) [41], with less patients experiencing severe adverse events. The identification of biomarkers of biological relevance supports the use of SHAP to identify predictive features.

A critical aspect of the SHAP analysis was the potential to define a numerical threshold for each feature value, whereby patients above or below this threshold transition from poor response to good response. The identification of numerical thresholds of M2 macrophages present an interesting case for further investigations. Due to the immunosuppressive nature of M2 macrophages, macrophage depletion therapies [42–44], such as anti-CSF1R [43], could be used in combination with ICI to boost favourable response. Several phase I clinical trials are investigating the effect of combining anti-CSF1R with different immunotherapies for solid cancers, including melanoma [45, 46]. Interestingly, our findings suggested that patients with M2 macrophage proportions exceeding approximately 30% had a higher likelihood of poor ICI response. In the future, when macrophage depletion therapy and their optimal dosage to combine with ICI are thoroughly investigated, it is possible that such a threshold could assist clinicians in determining whether adding macrophage depletion therapies to an ICI regimen would benefit melanoma patients.

While acknowledging the promise of explainable machine learning in selecting clinically relevant thresholds that may be useful for decision-making, it is important to emphasise that our findings are exploratory. Multiple lines of validation would need to occur to validate the clinical utility. Firstly, the numerical thresholds suggested by SHAP need to be compared against results generated by experimental methods, such as immunohistochemistry on larger patient cohorts. For example, immunohistochemistry is an established method for measuring CSF1R protein expression in melanoma patients [43], and could be used to validate levels of M2 macrophages in patients with good or poor response to ICI in future studies. Other explainability methods such as LIME [47] and Anchors [48], which are also model-agnostic, could be used to validate the features thresholds. Notably, LIME [47] has been utilised in cancer research in the classification of melanoma [49]. These validation studies will be crucial in translating our computational findings into personalised treatment strategies for melanoma patients.

In conclusion, explainable machine learning that integrates multimodal data can predict patient response to ICI, identify discriminatory features and suggest thresholds for their use. As additional ICI agents become available more treatment combinations will be considered. Future studies powered to predict patient response to different combinations of therapy will identify response biomarkers, but may also reveal candidate treatment targets and allow the personalisation of ICI treatment.

## DECLARATIONS

### Ethics approval and consent to participate

No patients were consented directly to this study. No participants were recruited for this study, the study included data from previously published studies. The work in this study was approved by the QIMR Berghofer human ethics research committee within project number P2095.

### Availability of data and materials

This project uses previously published DNA and RNA-Seq data. Data for Newell et al. [10] is available in the European Genome-phenome Archive (EGA) under study accession EGAS00001001552 [https://ega-archive.org/studies/EGAS00001001552] with dataset ID EGAD00001004869 [https://the.test.ega-archive.org/datasets/EGAD00001004869]. Data from Rozeman et al. [18] is available in the EGA with exome sequencing data under accession number EGAS00001004832 [https://ega-archive.org/studies/EGAS00001004832] and RNA-Seq data under EGAS00001004833 [https://ega-archive.org/studies/EGAS00001004833]. Sequence data for Riaz et al. [17] is available at the European Nucleotide Archive (ENA) database with ID’s PRJNA359359 and PRJNA356761. Data for Hugo et al. [19] is available at the ENA under accession PRJNA312948. Data reported in Campbell et al. [16] is available in the ENA under accession number PRJNA923698 for WES data and PRJNA923698 for RNA-Seq data. The R and Python code used for analyses in this study is available at https://github.com/MedicalGenomicsLab/multi-omics-ici-prediciton. Picard MarkDuplicates (v2.8.15) was accessed at https://broadinstitute.github.io/picard/. qCoverage and qSNP are available at https://github.com/AdamaJava/adamajava. SigProfilerExtractor was accessed at https://github.com/AlexandrovLab/SigProfilerExtractor. CIBERSORTx was accessed at https://cibersortx.stanford.edu/. TIDE was accessed at https://tide.dfci.harvard.edu/. All other data are available in the main text or the supplementary materials.

### Funding

National Health and Medical Research Council of Australia (NHMRC) Senior Research Fellowship APP1139071 (NW)

NHMRC Investigator Grant 2018244 (NW)

NHMRC Emerging Leader 1 Investigator Grant APP2008631 (OK)

The Mark Foundation for Cancer Research RG95043 (MCO)

Cancer Research UK Cambridge Centre CTRQQR-2021\100012 (MCO)

Academy of Medical Sciences SBF008\1170 (MCO)

NIHR i4i G121891 (MCO)

Australian Government as a Cooperative Research Centres Project (CRC P) (JVP, NW)

Australian Cancer Research Foundation (ACRF) – The ACRF Centre for Optimised Cancer Therapy (OK, JVP, NW)

### Author contributions

Conceptualization: JVP, NW

Methodology: VA, KAT, OK, JVP, NW

Software: KAT

Formal analysis: VA, KAT, JZ, NW

Investigation: VA, KAT

Data Curation: VA, KAT, LTK, JVP

Resources: SW, CL, LLH, CUB

Supervision: EDW, OK, JVP, NW

Project administration: JVP, NW

Funding acquisition: JVP, NW

Visualization: KAT, VA

Writing – Original Draft: VA, KAT, NW

Writing - Review & Editing: LTK, JZ, SW, CL, LLH, CUB, MCO, EDW, OK, JVP

## Acknowledgments

We are grateful to Sharon Hoyte, QIMR Berghofer bioinformatics, for her support in RNA-seq metrics. We wish to acknowledge helpful discussions with members of Max Kelsen Pty Ltd. This work and this research were performed on QIMR Berghofer computing infrastructure supported by the Australian Cancer Research Foundation (ACRF), The Ian Potter Foundation and The John Thomas Wilson Endowment. KT was the recipient of the Maureen and Barry Stevenson PhD Scholarship, we are grateful to Maureen Stevenson for her support.

