## Supplementary Figures for "Explainable machine learning identifies features and thresholds predictive of immunotherapy response"

**List of Supplementary Figures**

Fig. S1: Clinical, DNA and RNA in the test dataset

Fig. S2: Clinical and cancer intrinsic features significantly associated with immune checkpoint inhibitor response in the training data

Fig. S3: Cancer extrinsic features significantly associated with immune checkpoint inhibitor response in the training data

Fig. S4: Identification of features multi-collinearity and model-specific feature selection across different train-test permutations

Fig. S5: Training and test performance of single- and multi-omic machine learning models across different train-test permutations

Fig. S6: Performance of multi-omic Random Forest model with an independent test data

Fig. S7: Performance of multi-omic Random Forest on non-cutaneous melanoma

Fig. S8: SHAP scores of common features across all train-test permutations

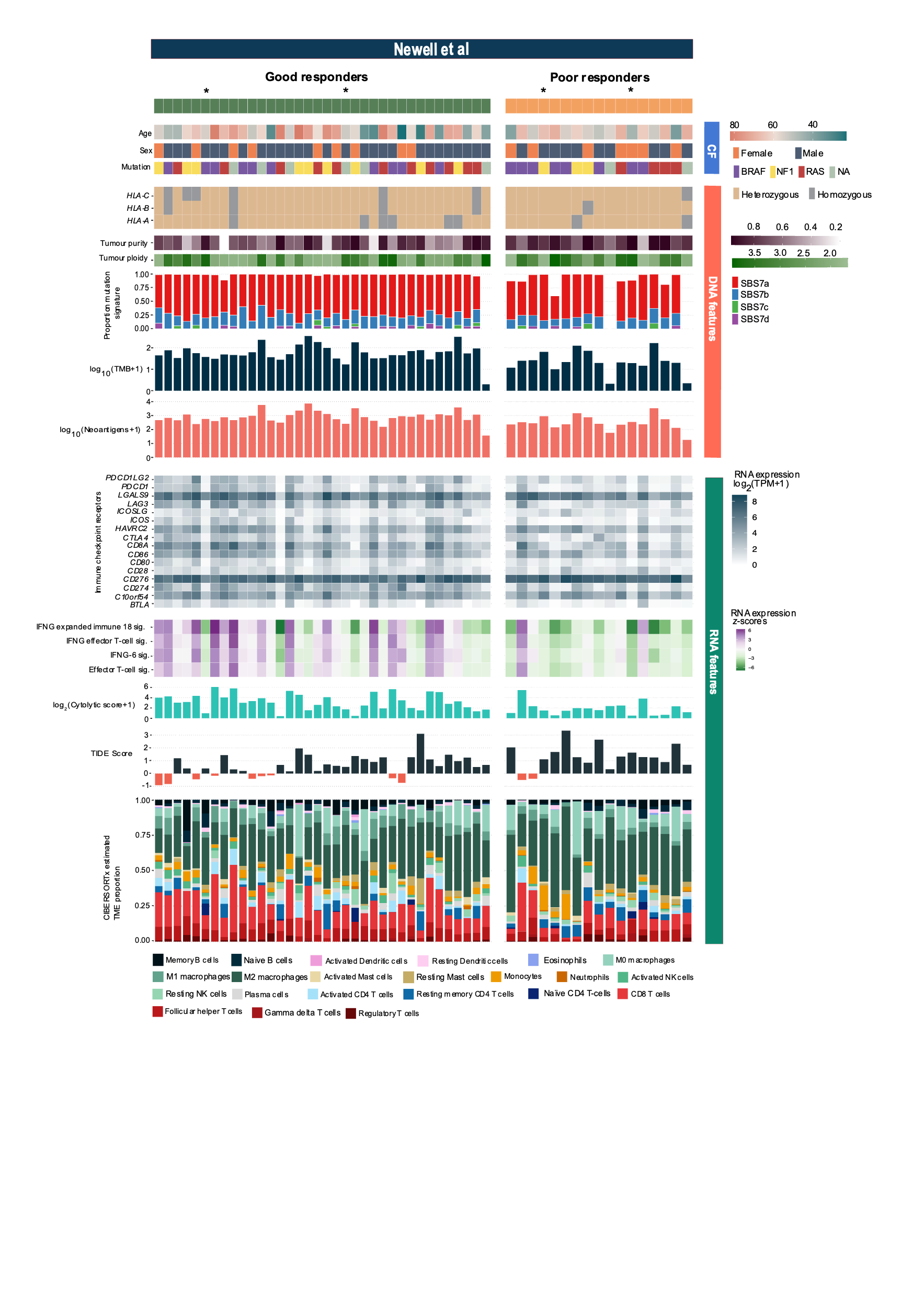

Fig. S1: Clinical, DNA and RNA in the test dataset. Overview of features from 53 patients sorted by Good (n= 36 green) vs Poor (n=17 orange) responders in the test dataset from Newell et al., with * indicating patients with stable disease. The clinical feature include: age (in years), sex (female, male) and mutation status of *BRAF*, *K/N/RAS* and *NF1* (NA indicates samples that have no identifiable mutation in these genes). Cancer intrinsic features determined by DNA sequence analysis are: zygosity of HLA allele, tumour purity (as a proportion of cancer cells), tumour ploidy, proportion of ultra-violet light mutation signatures (SBS7a, SBS7b, SBS7c, SBS7d), log transformed neoantigen load and log10 transformed TMB. Cancer extrinsic features determined by RNA sequence analysis are: log transformed TPM + 1 expression of selected immune checkpoint receptors, gene set signatures scores of Effector T-cell, IFNg-6, a combined IFNg/effector T cell and a combined 18-gene expanded IFNg signatures, as well as log­_2_-transformed cytolytic score and TIDE score and the proportion of immune cells estimated by deconvolution of RNA-seq data by CIBERSORTx. CF: clinical features, HLA: human leukocyte antigen, IFNg: interferon gamma, SBS: single base substitution, TIDE: tumour immune dysfunction and exclusion, TMB: tumour mutation burden, TME: tumour microenvironment.

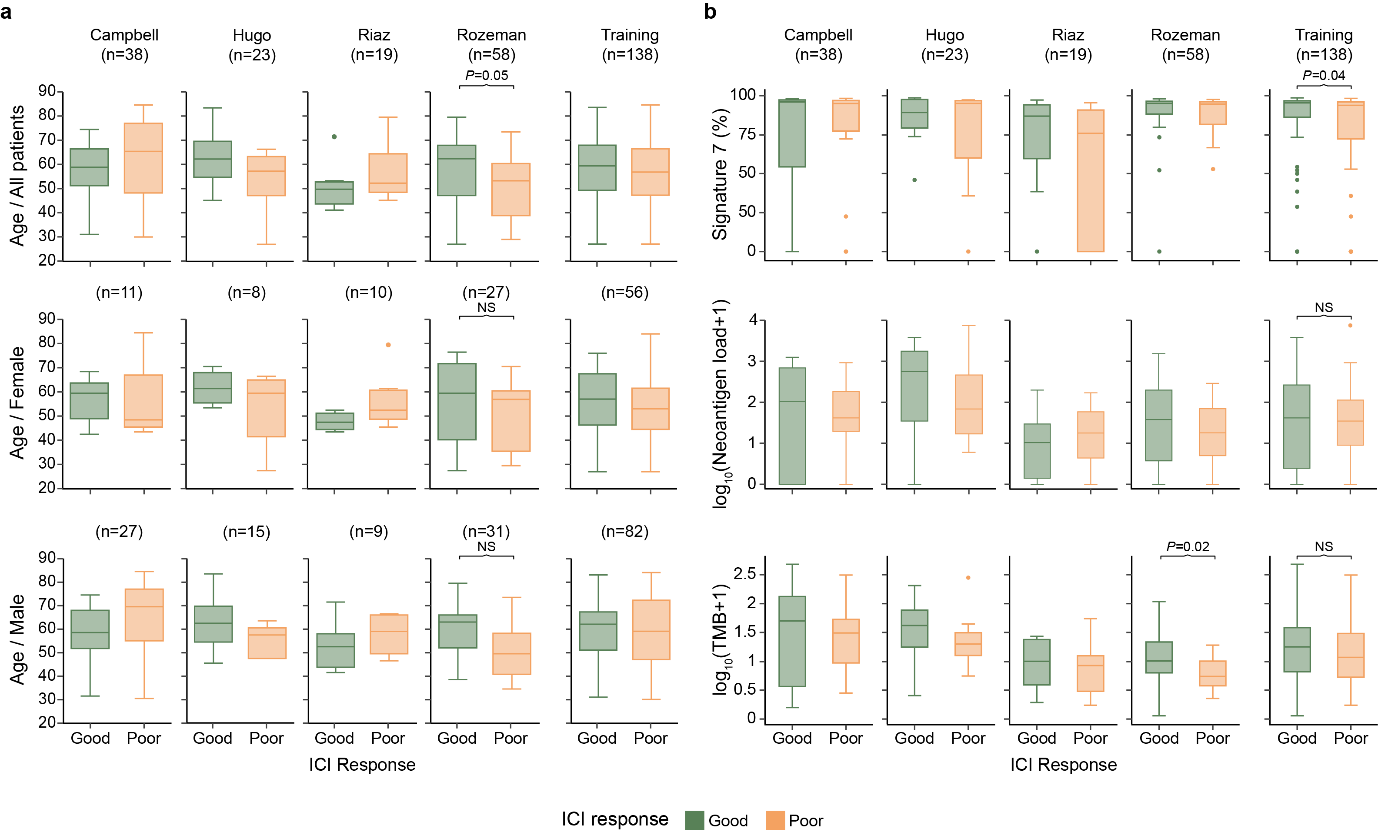

**Fig. S2: Clinical and cancer intrinsic features significantly associated with immune checkpoint inhibitor response in the training data.** **A**) Box plots showing the age of patients in years (y-axis) with data grouped into patient response to ICI with good (green) and poor (orange). The patients are aggregated by biological sex (all patients upper plot, female middle plots, male lower plots). **B**) Box plots showing the cancer intrinsic feature values (y-axis) with data grouped into patient response to ICI with good (green) and poor (orange). The three cancer intrinsic features are Signature 7 shown as a proportion, log_10_(neoantigen load +1) and log_10_(TMB+1). For all box plots the boxplot depicts the middle 50% of data. The whiskers extend to the maximum and minimum values, not including outliers. Outliers are values that deviate more than 1.5 times the interquartile range from either Q1 or Q3. In each panel the boxplots are aggregated into five plots with the first four containing good versus poor ICI response for each individual training dataset (Campbell n=38, Hugo n=23, Riaz n=19 and Rozeman n=58), and the last plot containing the good versus poor ICI response for all training data (n=138). Boxplots with significant p-values ($\boldsymbol{\leq0.05}$, MWU) between good and poor responders are annotated with the corresponding p-value. ICI: immune checkpoint inhibitor, MWU: Mann-Whitney U-test, NS: not significant, TMB: tumour mutation burden.

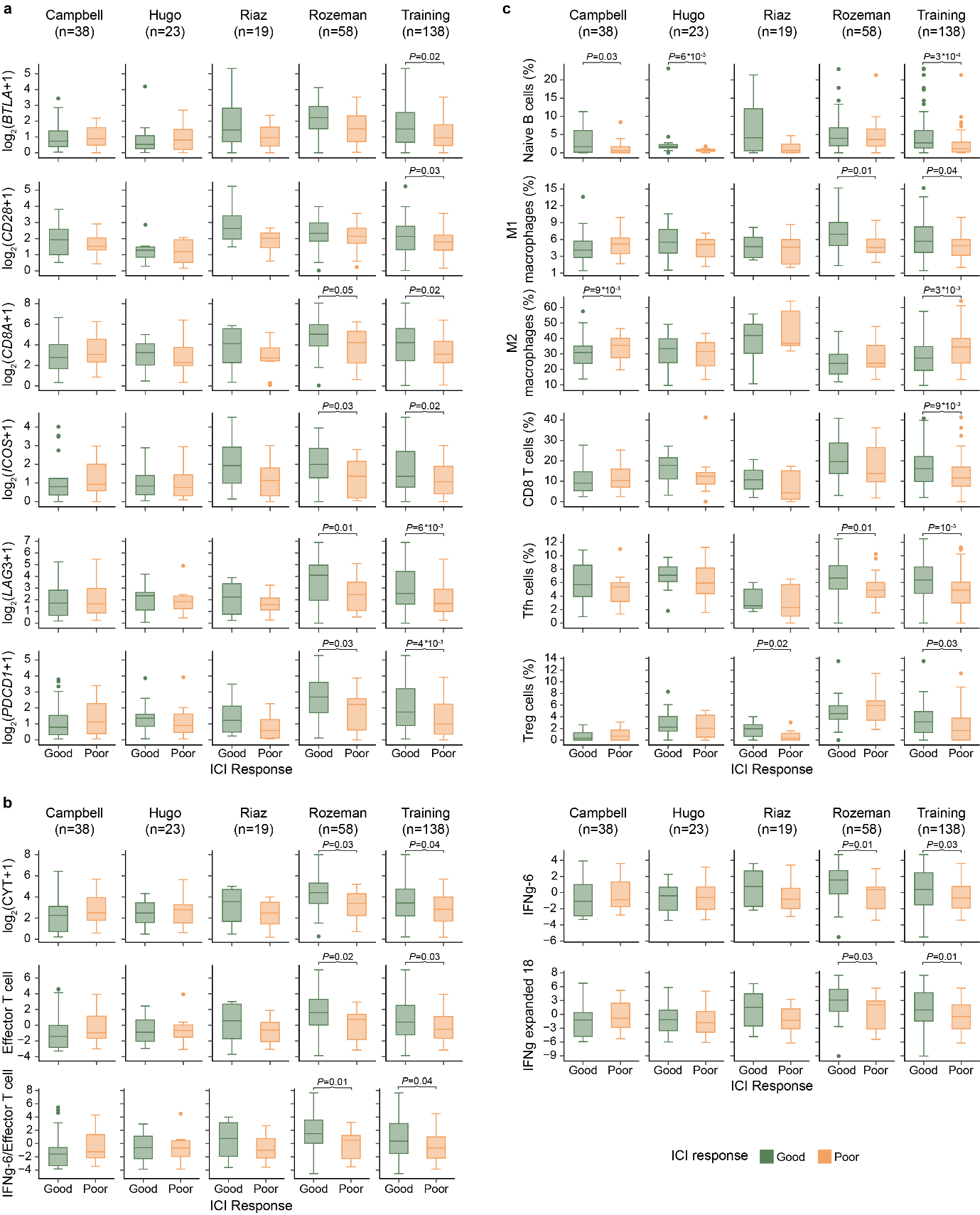

Fig. S3: Cancer extrinsic features significantly associated with immune checkpoint inhibitor response in the training data. A) Box plots showing the expression (log_2_(TPM + 1)) of six immune checkpoint receptor marker genes (y-axis) with data grouped into patient response to ICI with good (green) and poor (orange). B) Box plots showing log2p1-transformed values of cytolytic scores and non-transformed values of Effector T-cell, IFNg-6, IFNg/effector T cell and IFNg expanded 18 signatures (y-axis) with data grouped into patient response to ICI with good (green) and poor (orange). C) Box plots showing the CIBERSORTx-estimated proportions of immune cell types (y-axis) with data grouped into patient response to ICI with good (green) and poor (orange). Each boxplot depicts the middle 50% of data. The whiskers extend to the maximum and minimum values, not including outliers. Outliers are values that deviate more than 1.5 times the interquartile range from either Q1 or Q3. In each panel the boxplots are aggregated into five plots with the first four containing good versus poor ICI response for each individual training dataset (Campbell n=38, Hugo n=23, Riaz n=19 and Rozeman n=58), and the last plot containing the good versus poor ICI response for all training data (n=138). Boxplots with significant p-values ($\boldsymbol{\leq0.05}$, Mann-Whitney U-test) between good and poor responders are annotated with the corresponding p-value. CYT: cytolytic, ICI: immune checkpoint inhibitor, IFNg: interferon gamma, log2p1: logarithm base 2 of value plus 1, NK cells: Natural Killer cells, Tfh: follicular helper T cells, Treg cells: regulatory T cells.

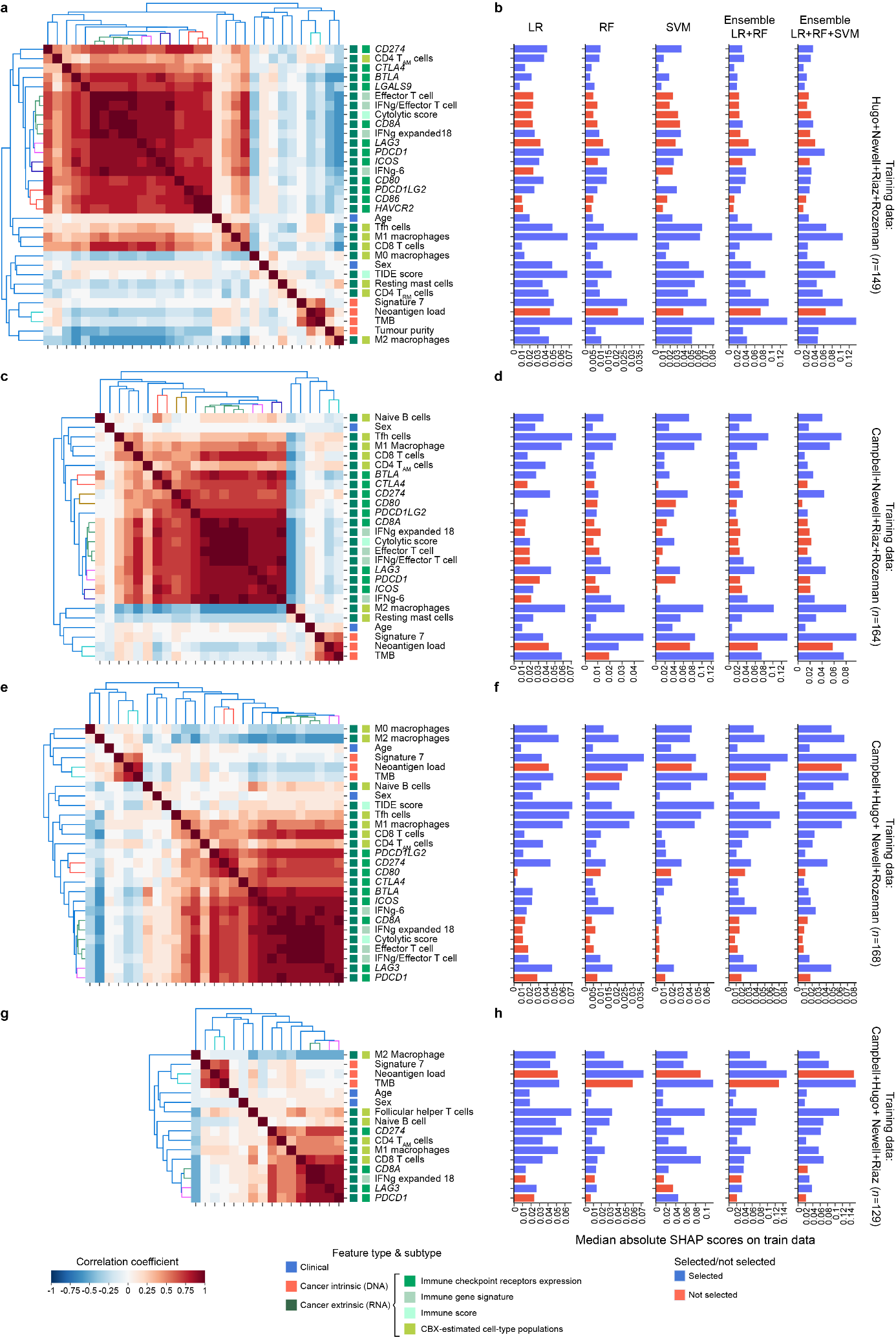

**Fig. S4: Identification of features multi-collinearity and model-specific feature selection across different train-test permutations.** Feature collinearity analysis and SHAP-based feature selection for when **(A-B)** Campbell et al. was used as test data (thus training data include cutaneous and non-stable-disease samples from Hugo et al, Newell et al., Riaz et al. and Rozeman et al. (n=149)), **(C-D)** Hugo et al. was used as test data (thus training data include cutaneous and non-stable-disease samples from Campbell et al, Newell et al., Riaz et al. and Rozeman et al. (n=164)), **(E-F)** Riaz et al. was used as test data (thus training data include cutaneous and non-stable-disease samples from Campbell et al., Hugo et al., Newell et al. and Rozeman et al. (n=168)), **(G-H)** Rozeman et al. was used as test data (thus training data include cutaneous and non-stable-disease samples from Campbell et al., Hugo et al., Newell et al. and Riaz et al. (n=129)). In **(A)**, **(C)**, **(E)**, and **(G)**, correlation plots showing pair-wise Pearson’s r coefficient between pairs of numerical-numerical variables, and Eta Correlation Ratio between pairs of numerical-categorical variables. The correlation scale is depicted beneath the plots with darker shade of blue representing stronger negative correlation, and darker shade of red representing stronger positive correlation. The feature types are coloured by clinical: blue, Cancer intrinsic (DNA): orange and Cancer extrinsic (RNA): green. Cancer extrinsic are further separated into expression of immune checkpoint receptor marker genes, immune signature scores and the CBX-estimated proportions of immune cell types. For each hierarchical clustering output accompanying the pairwise correlation matrix, different sets of highly correlated features are indicated by coloured clusters. In **(B)**, **(D)**, **(F)**, and **(H)**, median absolute SHAP scores (x-axis) computed over training samples for each feature (y-axis) after training each model on the feature set. For each model, blue bars represent selected features, and red bars show features discarded among the four highly correlated feature clusters. CBX: CIBERSORTx, CD4 T_AM_ cells: activated resting memory CD4 T cells, CD4 T_RM_ cells: resting memory CD4 T cells, IFNg: interferon gamma, LR: logistic regression, RF: random forest, SVM: support vector machine, Tfh cells: follicular helper T cells, TIDE: T-cell immune dysfunction and exclusion, TMB: tumour mutation burden, Treg cells: regulatory T cell.

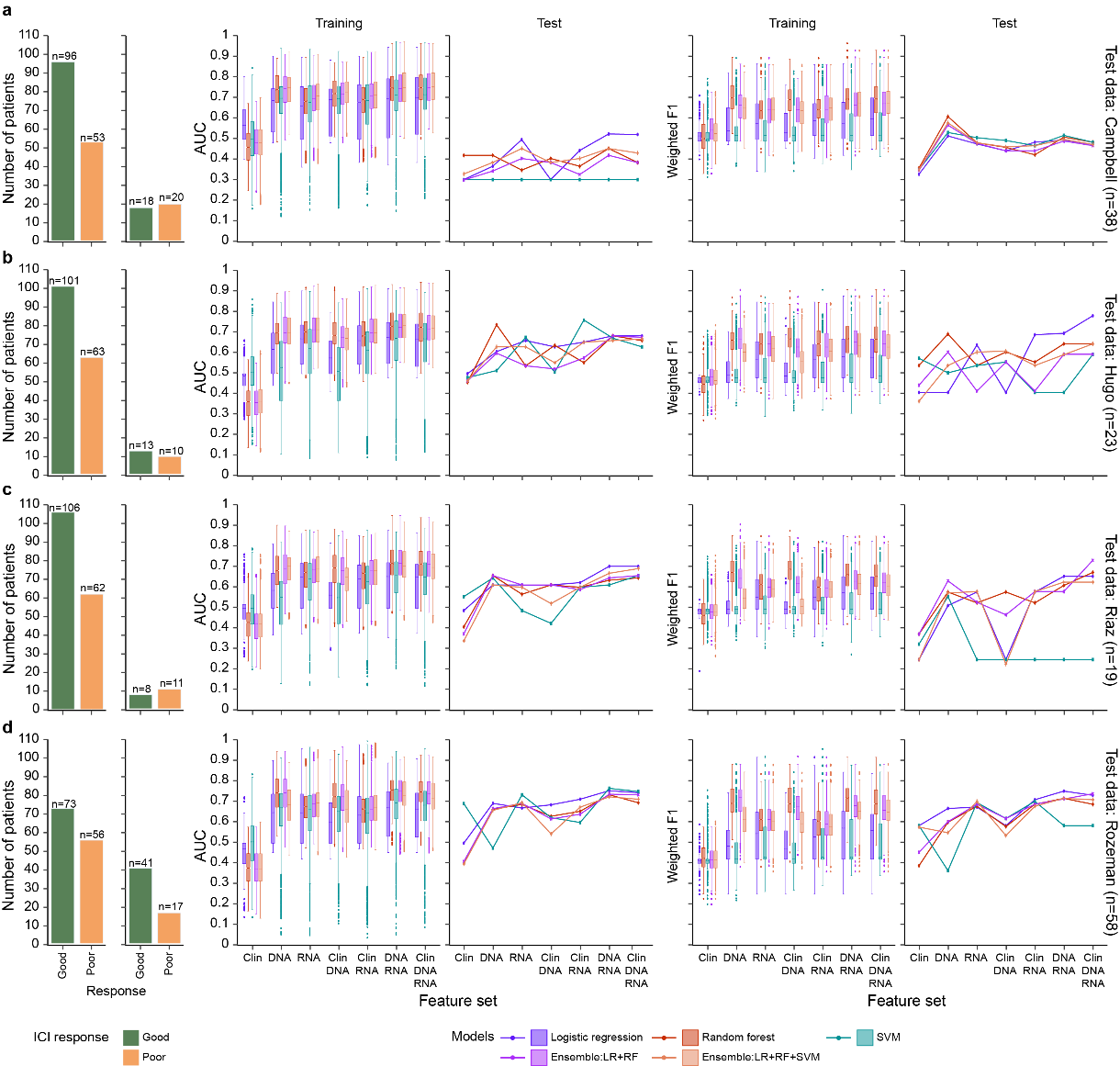

Fig. S5: Training and test performance of single- and multi-omic machine learning models across different train-test permutations. AUC-ROC and Weighted F1 values achieved by the machine learning models when cutaneous and non-stable-disease samples from (A) Campbell et al. (n=38), (B) Hugo et al. (n=23), (C) Riaz et al. or (D) Rozeman et al. were used as test data. Performance metrics for each train-test permutation are accompanied by bar plots (left-most column) representing number of good (green) and poor (orange) responders in the corresponding training data. Box plots of AUC-ROC and weighted F1 values (y-axis) represent models’ performance achieved during cross-validation within the corresponding training data. Each boxplot depicts the middle 50% of F1 or AUC-ROC values, encompassing the first quartile (Q1), the median, and the third quartile (Q3). The whiskers extend to the maximum and minimum weighted F1/AUC-ROC values, not including outliers. Outliers are AUC-ROC/ weighted F1 values that deviate more than 1.5 times the interquartile range from either Q1 or Q3. Line plots represent single AUC-ROC and weighted F1 scores (y-axis) of the best models, based on training performance, on the corresponding test data. For all boxplots and line plots, higher AUC-ROC/weighted F1 values indicate better performance. AUC-ROC: area under the curve of the receiver operating characteristic curve, LR: logistic regression, RF: random forest, SVM: support vector machine.

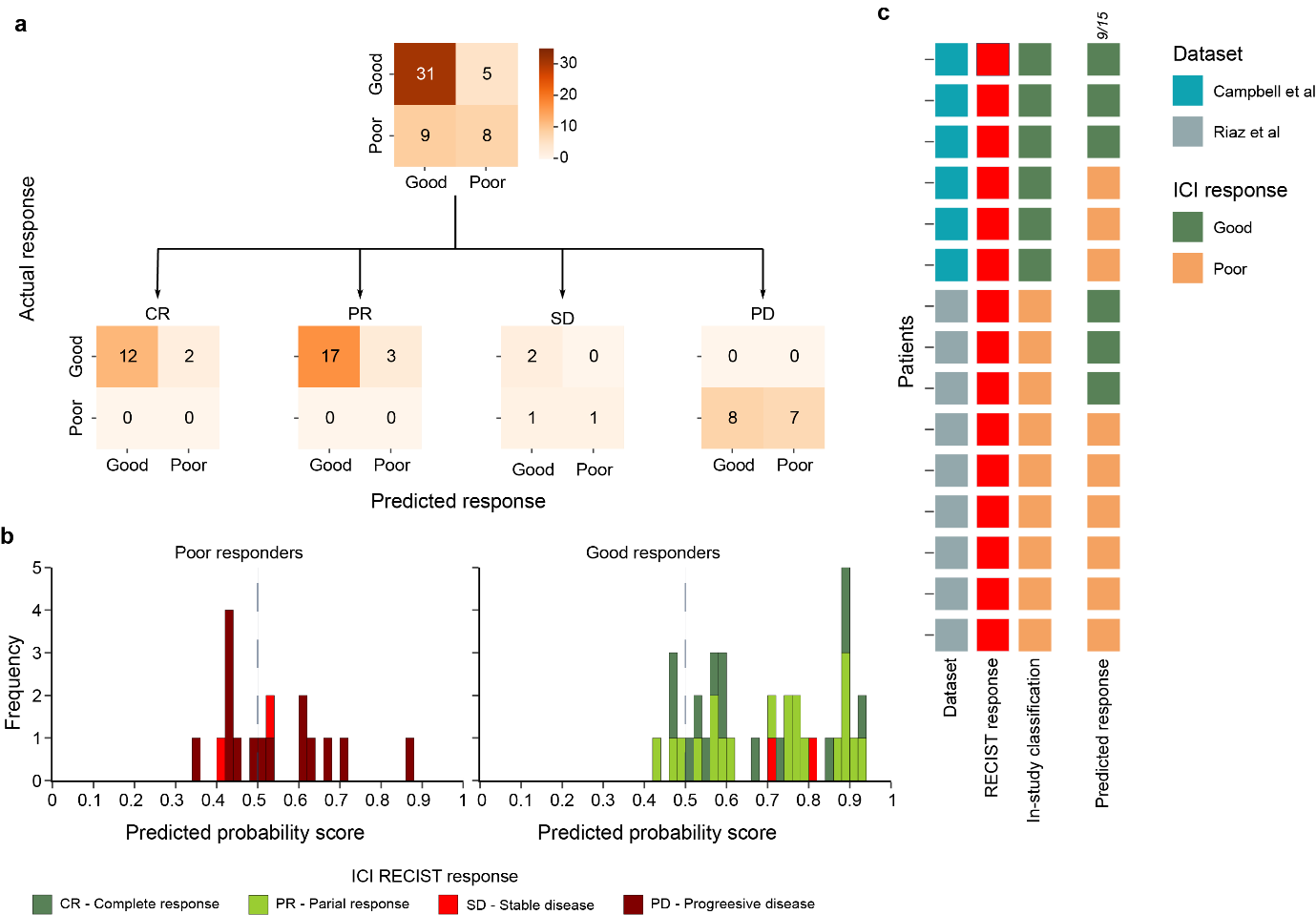

Fig. S6: Performance of multi-omic Random Forest model with an independent test data. Evaluation of results produced by the Random Forest (RF) model trained using DNA and RNA features on 53 test samples from Newell et al. A) Confusion matrices of the model’s predictions of Good and Poor, broken down into RECIST response (CR, PR, SD, PD) in the second row. B) The frequency (y-axis) of probability scores (x-axis), color-coded by RECIST response (CR: teal, PR: green, SD: red, PD: dark red). The vertical line at 0.5 dictates the good versus poor response threshold. Predictions to the left of this line were classified as poor response, while predictions to the right were classified as good response. C) Predicted responses of patient with stable disease from Campbell et al. (n=6) and Riaz et al. (n=9). CR: Complete response; PR: Partial response, SD: Stable disease, PD: Progressive disease.

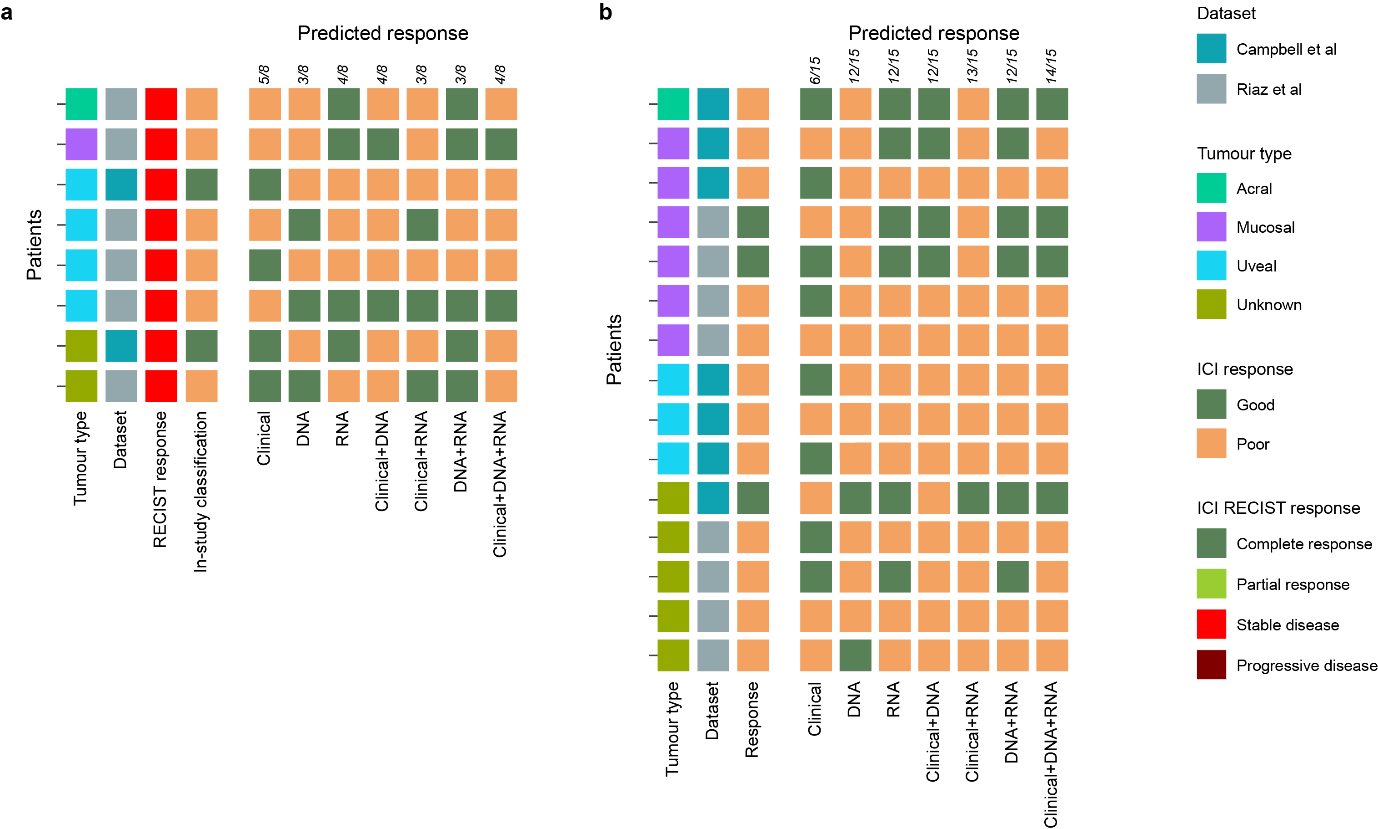

Fig. S7: Performance of multi-omic Random Forest on non-cutaneous melanoma. Predictions of ICI response for 23 non-cutaneous test samples made by the Random Forest (RF) models trained on clinical features, DNA features, RNA features, or both Clinical and DNA features, or both Clinical and RNA features, or DNA and RNA features, or all features (Clinical, DNA and RNA). Models’ predictions are aggregated by, A) patients with stable disease (n=8) and B) patients with a good or poor response (n=15). Predictions of each model are accompanied by the number of correct prediction (x/6 or x/15). Response shown is from the original study (Campbell et al. or Riaz et al.), predicted responses (Good or Poor) are shown for each model. For each panel, patient identifiers are on the left, with features of these samples shown in the left columns including tumour type (acral: green, mucosal: purple, unknown: green and uveal: blue), dataset (Campbell et al.: teal and Riaz et al.: grey) and the actual ICI response for each patient (Good: green and Poor: orange).

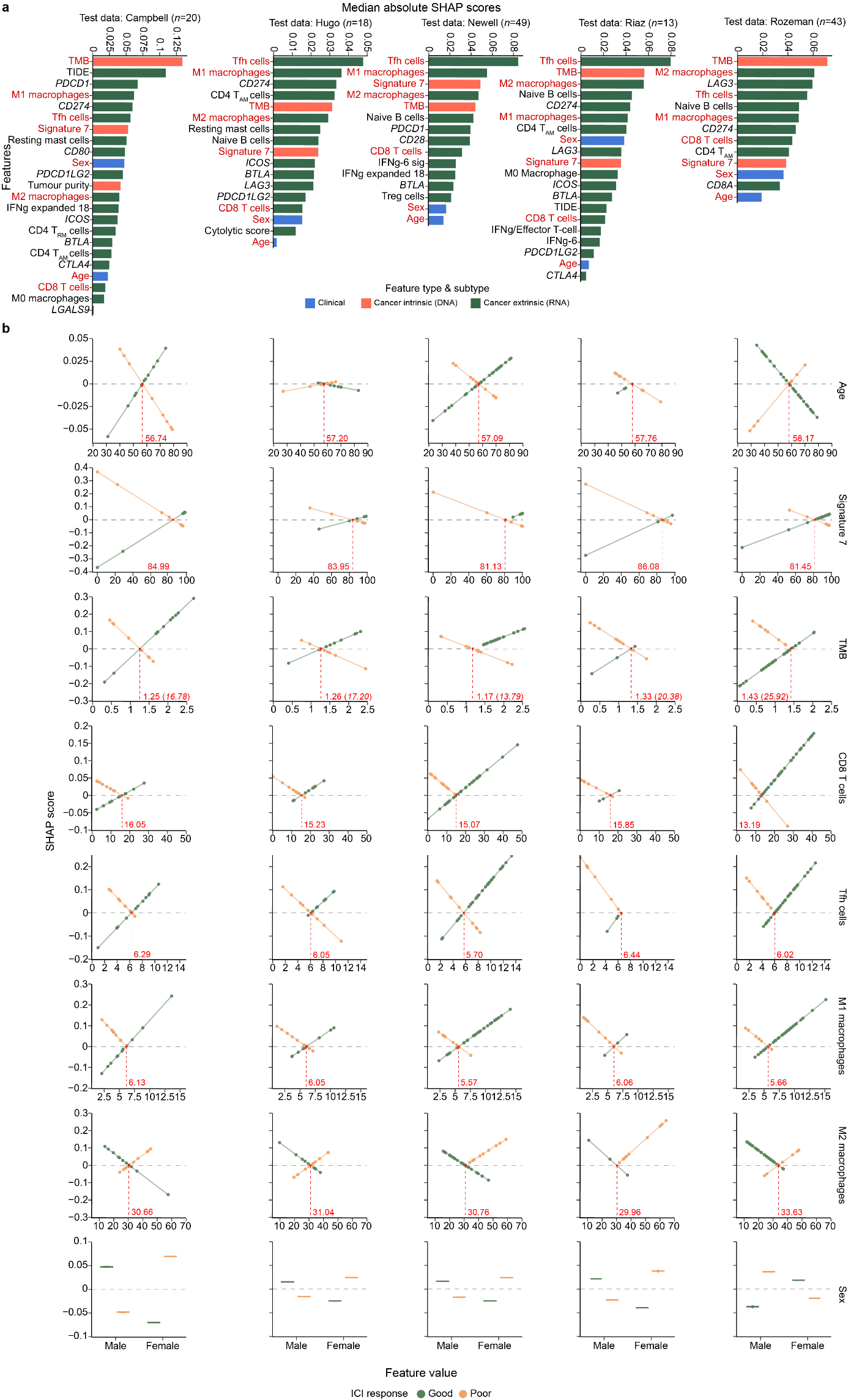

Fig. S8: SHAP scores of common features across all train-test permutations. Bar plots illustrating median absolute SHAP scores (A) and scatter plot illustrating raw SHAP scores against feature values (B) computed from correctly predictions by the multi-omic (Clinical+DNA+RNA) LR model when cutaneous and non-stable-disease samples from Campbell et al. (first column, n=20), Hugo et al. (second column, n=18), Newell et al. (third column, n=49), Riaz et al. (fourth column, n=13) or Rozeman et al. (last column, n=43) were used as test data. In (a), bar plots were coloured by feature types, and eight common features included in all train-test permutations (Age, Signature 7, TMB, CD8 T cells, Follicular helper T cells, M1 macrophages, M2 macrophages and Sex) were highlighted in red. The direction of influence (- or + SHAP scores) of each feature is not captured. In b), only SHAP scores and feature values of the eight common features are displayed, with each data point representing a patient in the corresponding test data. The patients are coloured by response (good: green and poor: orange). CIBERSORTx-estimated cell-type proportions are depicted in percentages, and TMB values are depicted on log10p1 scale (accompanied by converted linear values in bracket). The horizontal dashed line represents the SHAP score of 0, indicating that a feature has no influence on a particular prediction. Green and orange lines represent best-fitted lines identified by Ordinary Least Squares (OLS) for good and poor responders, respectively. Red vertical lines represent manually annotated intersections between good and poor, assisted by intersections of good and poor OLS-fitted lines. CD4 T_AM_ cells: activated resting memory CD4 T cells, CD4 T_RM_ cells: resting memory CD4 T cells, ICI: immune checkpoint inhibitor, IFNg: interferon gamma, SHAP: SHapley Additive exPlanations, Tfh cells: follicular helper T cells, TMB: Tumour Mutational Burden.
