## Supplementary Notes for "Explainable machine learning identifies features and thresholds predictive of immunotherapy response"

**List of Supplementary Notes**

Supplementary Note 1: Feature generation

Supplementary Note 2: Multi-omic machine learning workflow

### Supplementary Note 1: Feature generation

The DNA and RNA sequence data from 5 published studies [1-5] was accessed in FASTQ format and re-analysed to ensure data uniformity. Using this data, we assembled a total of 57 features from each patient that were used to train and test the model. The features consisted of clinical, tumour DNA and tumour RNA features. The clinical features were patient age at diagnosis, patient biological sex and binary mutation status of *BRAF*, *KRAS/NRAS* or *NF1* (“Mutated” or “Not mutated”). The DNA features were selected to capture cancer intrinsic features that may be predictive of ICI response [6] and included tumour HLA zygosity, TMB, neoantigen load, tumour purity, tumour ploidy, and mutational signature 7 linked to UV light and sun exposure that have been previously described in melanoma [7, 8]. Features from RNA-Seq data were selected to represent cancer extrinsic properties of the TME that may be indicative of ICI response [6]. These features included the gene expression level of 16 immune checkpoint receptors, four immune gene signatures previously reported to be associated with ICI response [9], two immune scores (TIDE [10] and the cytolytic score [11]), as well as the relative proportion of 22 immune cells within the TME measured by CIBERSORTx [12]. All features used in this study are defined in **Supplementary Materials Table 1**.

**Supplementary Materials Table 1. Definitions of clinical, cancer intrinsic (DNA) and extrinsic (DNA) features used in this study**

| **Type** | **Subtype** | **Feature** | **Description** |
| --- | --- | --- | --- |
| Clinical | N/A | Age | Age of patient at start of treatment or diagnosis |
|  |  | Sex | Biological sex of patient at birth |
| DNA | N/A | Signature 7 | The proportion of mutational signatures that are assigned to Signature 7 – total (i.e. sum of SBS7a, SBS7b, SBS7c and SBS7d) |
|  |  | HLA-A Alleles | The HLA-A allele of the tumour |
|  |  | HLA-B Alleles | The HLA-B allele of the tumour |
|  |  | HLA-C Alleles | The HLA-C allele of the tumour |
|  |  | *BRAF* mutation status | Mutation status of BRAF (somatic mutations at V600) |
|  |  | *NF1* mutation status | Mutational status of NF1 (all somatic mutations) |
|  |  | *N/K-RAS* mutation status | Mutational status of RAS (somatic mutations at G12 or Q61) |
|  |  | Neoantigen load | The number of neoantigens in the tumour with Inhibitory Concentration (IC50) of ≤500nM. log10(x+1) |
|  |  | Tumour mutation burden | The total number of somatic non-synonymous mutations per Mb in the coding region. log10(x+1) |
|  |  | Tumour ploidy | The ploidy of the tumour |
|  |  | Tumour purity | The percentage of cancer cells within the tumour |
| RNA | Immune checkpoint receptors expression | *BTLA* | TPM counts as log2(x+1) |
|  |  | *C10orf54* |  |
|  |  | *CD274* |  |
|  |  | *CD276* |  |
|  |  | *CD28* |  |
|  |  | *CD80* |  |
|  |  | *CD86* |  |
|  |  | *CD8A* |  |
|  |  | *CTLA4* |  |
|  |  | *HAVCR2* |  |
|  |  | *ICOS* |  |
|  |  | *ICOSLG* |  |
|  |  | *LAG3* |  |
|  |  | *LGALS9* |  |
|  |  | *PDCD1* |  |
|  |  | *PDCD1LG2* |  |
|  | Immune gene signature | Effector T-cell | Immune signature score[13, 14] |
|  |  | IFNg-6 |  |
|  |  | IFNg/Effector T-cell |  |
|  |  | IFNg expanded 18 |  |
|  | Immune score | Cytolytic score | Immune signature score from the geometric mean of gene expression of *GZMA* and *PRF1*. log2(x+1) |
|  |  | TIDE score | Immune signature score - The Tumour Immune Dysfunction and Exclusion (TIDE) signature[15] |
|  | CBX-estimated cell-type population | Memory B cells | Relative proportion of 22 immune cell types in LM22 signature matrix. |
|  |  | Naïve B cells |  |
|  |  | Activated dendritic cells |  |
|  |  | Resting dendritic cells |  |
|  |  | Eosinophils |  |
|  |  | M0 macrophages |  |
|  |  | M1 macrophages |  |
|  |  | M2 macrophages |  |
|  |  | Activated mast cells |  |
|  |  | Resting mast cells |  |
|  |  | Monocytes |  |
|  |  | Neutrophils |  |
|  |  | Activated NK cells |  |
|  |  | Resting NK cells |  |
|  |  | Plasma cells |  |
|  |  | Activated memory CD4 T cells |  |
|  |  | Resting memory CD4 T cells |  |
|  |  | Naive CD4 T cells |  |
|  |  | T CD8 T cells |  |
|  |  | Follicular helper T (Tfh) cells |  |
|  |  | Gamma delta T cells |  |
|  |  | Regulatory T (Treg) cells |  |

*CBX: CIBERSORTx, HLA: human leukocyte antigen, NK: natural killer

#### DNA feature generation

#### *Somatic mutation analysis:*

DNA sequence files were adapter trimmed with (Cutadapt v1.9) and reads were mapped to human reference genome GRCh37 using BWA-MEM [16] (v0.7.15). Duplicate reads were marked with Picard MarkDuplicates (v2.8.15). The sequence read depth was estimated for tumour and normal samples using qCoverage (v0.7). The mean read depth for each study was: Campbell el al. [1] tumour samples 45.13x (range 9–67) and normal samples 48.7x (range 7–88); Rozeman et al. [5] tumour samples 117x (range 77–162) and normal samples 118x (range 84–163); Riaz et al. [4] tumour samples 58.5x (range 37–83) and normal samples 27.7x (range 17–47); Hugo et al. [2] tumour samples tumour samples 71.0x (range 43–129) and normal samples 70.5x (range 35–141) and Newell et al. [3] tumour samples 60.5x (range 31–80) and normal samples 25x (range 18–31) (**Table S3**). Somatic single nucleotide variants were identified using the consensus of two approaches, qSNP[17] (version 2.0) and GATK HaplotypeCaller[18] (version 3.3-0). SnpEff [19] was used to annotate variants with gene consequence. Tumour mutation burden (TMB) was defined as the number of non-synonymous somatic mutations (SNVs and indels) per megabase in the coding region of the genome. Mutation status of *BRAF*, *KRAS*, *NRAS* and *NF1* was determined by looking for the presence mutations at specific hotspot positions of *BRAF* (V600), *KRAS* and *NRAS* (G12 or Q61) and all non-synonymous mutations in *NF1*. Tumour purity and ploidy was assessed using ascatNGS [20] for WGS data and sequenza for exome sequence data.

#### *Mutational signatures*

We applied SigProfilerExtractor (v1.1.20) to somatic single nucleotide mutations from all samples within the five datasets [1-5]. In brief, SigProfilerExtractor implements nonnegative matrix factorisation (NMF) [21, 22] to find a set of single base substitution (SBS) signatures ranging from 2 to 12 for each sample using 100 iterations. The extracted signatures from optimal solution with rank=3 was further decomposed to COSMIC SBS signatures (v3.3). To prevent overfitting, signatures contributing less than 10% to a sample were excluded. Finally, we collected signature proportions of ultraviolet (UV) light exposure related signatures (SBS7a, SBS7b, SBS7c, SBS7d) as the UV signature has been reported as dominant in the majority of melanoma patients [7].

#### *HLA typing and neoantigen prediction*

Class I HLA genotypes were determined for paired tumour-control whole-genome and exome datasets using Optitype [23] (v1.3.1) with default parameters, based on HLA genotypes we manually determined whether an individual was homozygous or heterozygous. The pVAC-Seq[24] (v2.0.4) pipeline, was executed with default parameters to predict neoantigens from somatic single variant mutations, and NetMHCpan (v4.0) was employed to estimate binding affinity. Mutations were annotated for wildtype and mutant peptide sequences using the ENSEMBL Variant Effect Predictor [25] (v86). Epitopes with an Inhibitory Concentration (IC50) ≤500nM were deemed as potential neoantigens binding to HLA alleles and were used to calculate the neoantigen load.

#### RNA feature generation

#### *RNA-Seq alignment and gene counts*

RNA sequencing datasets were aligned to the human GRCh38 using STAR aligner [26] (v2.5.2a). Sequencing adapters were trimmed using Cutadapt [27] (v1.9). Gene annotation, transcript and exon features of Gencode GRCh38 (release 22) were used to compute counts of individual genes for all RNA-Seq samples. Quality metrics were assessed using RNA-SeQC [28] (v1.1.8) (**Table S4**). Gene expression was estimated using RSEM (v1.2.30) [29]. According to the counts, gene annotations and sequencing depth of the samples, transcripts-per-million (TPM) values were computed across samples.

#### *Immune receptors expression, immune scores and gene signatures*

The log_10_(TPM+1) values of 16 genes encoding immune checkpoint receptors *BTLA, C10orf54, CD274, CD276, CD28, CD80, CD86, CD8A, CTLA4, HAVCR2, ICOS, ICOSLG, LAG3, LGALS9, PDCD1, PDCD1LG2* was applied to estimate the gene expression across all the samples. The cytolytic score[11] was calculated as the geometric mean of gene expression of *GZMA* and *PRF1* (using TPM, 0.01 offset). The Tumour Immune Dysfunction and Exclusion (TIDE) signature [10] was calculated using TPM values transformed to log_2_(TPM +1) and then normalised using the recommended method. The gene signatures analysed were a six gene IFNγ signature (IFNγ-6), a related 18 gene expanded immune signature (IFNy expanded immune 18) [9], an effector T cell signature [13], a combined IFNy/Effector T-cell signature [30]. For the IFNy-6, IFNy expanded immune 18, Effector T-cell, IFNy/Effector T-cell log_2_(TPM counts +1) were used and a single gene set score for each gene set and patient was calculated using the z-score method in the GSVA R package [31].

#### *TME deconvolution*

The deconvolution of immune cells within the tumour microenvironment was conducted using CIBERSORTx [12] using TPM counts from RSEM using 500 permutations, B-mode batch correction, and the provided LM22 gene signature file comprising 22 immune genes with quantile normalisation disabled. Of note, while the immune scores and gene signatures were generated by applying the related methods on all RNA-Seq samples, CIBERSORTx was run separately for RNA-Seq samples from each dataset. This was to comply with CIBERSORTx execution instructions (Item 9, Section 4. Notes), which recommended that CIBERSORTx be run separately for mixtures from different datasets with known batch effects.

### Programming language and libraries

#### Python

We developed the machine learning models in this study using Python v3.9.17. The machine learning pipelines were built using a combination of scikit-learn v1.3.2, scikit-bio v0.5.9, pandas v2.2.2, numpy v1.24.4, scipy v1.10.1, and openpyxl v3.1.2. Where necessary, data visualisation was done using the Python libraries plotly v5.16.1, matplotlib v3.8.3 and seaborn 0.13.0. Features importance analysis was done using shap v0.45.0. Other supporting libraries were orca v1.8, kaleido v0.2.1, click v8.1.7, and tqdm 4.65.0. Additionally, running the interactive Jupyter notebooks also requires nb-conda v2.2.1 and ipykernel 5.5.5.

## R

R version used in this project was 4.3.1. The required packages are tidyverse v2.0.0, GSVA v 01.5.0.

### Supplementary Note 2: Multi-omic machine learning workflow

#### Soft voting in ensemble machine learning

We built two ensemble machine learning models in this study, with one model consisting of Logistic Regression (LR) and Random Forest (RF) (i.e. Ensemble:LR+RF), and another model consisting of LR, RF and SVM (Ensemble:LR+RF+SVM). We used soft voting to reconcile predictions of component models for both ensembles. This means probability scores of component models are averaged to produce the final probability score of each ensemble. More specifically, for the Ensemble:LR+RF model:

(1) ${Prob}_{Ensemble:LR+RF}\left( Good response \right)= \frac{1}{2} \left[ {Prob}_{LR}( )+{Prob}_{RF}( ) \right]$

and for the Ensemble:LR+RF+SVM model:

(2) ${Prob}_{Ensemble:LR+RF+SVM}\left( Good response \right)=\frac{1}{3} \left[ {Prob}_{LR}( )+{Prob}_{RF}( )+{Prob}_{SVM}( ) \right]$

After this averaging step, each ensemble’s predictions are decided by a thresholding function:

(3)${Pred}_{Ensemble}=\left\{ \begin{aligned} \text{"poor response"} if {Prob}_{Ensemble}<0.5 \\ \text{"good response"} if {Prob}_{Ensemble}\geq0.5 \end{aligned} \right.$

We note that an alternative to soft voting is hard voting, whereby the final prediction of an ensemble is decided as the most common prediction among the component models (i..e. majority rule). However, this approach would have not allowed us to retrieve probability scores of each ensemble, and we therefore chose soft voting.

#### Data transformation

We use the machine learning scikit-learn function $Pipeline()$ to transform input data into each machine learning model. Categorical features were one-hot-encoded using the function $OneHotEncoder(handle\_unknown="ignore")$. On the other hand, we used the function $StandardScaler()$ to scale numerical features to unit variance, resulting in all features centered around 0 and having a variance of 1. This technique is otherwise known as z-score scaling. When no categorical feature or numerical feature is present in the input data, e.g. all RNA features are numerical, the corresponding transformation step is skipped.

#### Model training and hyperparameter optimisation

Supplementary Materials Table 2 lists optimised hyperparameters and the values we searched through for each hyperparameter. Supplementary Materials Table 3 lists all non-optimised hyperparameters and their fixed values.

**Supplementary Materials Table 2. List of optimised hyperparameters for each machine learning model**

| **Classifier** | | **Hyperparameter** | **Searched values** |
| --- | --- | --- | --- |
| LR | | max_iter | [10, 100, 1000] |
|  |  | C | [0.0001, 0.005, 0.001, 0.01, 0.5, 1.0, 10.0] |
|  |  | l1_ratio | [0.1 0.2 0.3 0.4 0.5 0.6 0.7 0.8 0.9] |
|  |  | penalty | ['l1', 'l2', 'elasticnet'] |
|  |  | solver | ['liblinear', 'lbfgs', 'newton-cg', 'saga'] |
| RF | | n_estimators | [1000, 2000, 5000] |
|  |  | max_depth | [6, 8, 10] |
|  |  | criterion | ['gini', 'entropy'] |
|  |  | max_features | ['sqrt', 0.5, 0.75] |
| SVM | | kernel | ['linear', 'rbf', 'sigmoid'] |
|  |  | C | [0.0001, 0.001, 0.01, 0.1, 1, 10.0, 100.0] |
|  |  | degree | [2, 3, 4, 5] |
|  |  | gamma | [1e-06, 1e-05, 0.0001, 0.001, 0.01, 0.1, 1, 10.0, 100.0, 1000.0] |
| Ensemble:LR+RF | LR | max_iter | [100, 1000] |
|  |  | C | [0.0001, 0.001, 0.01, 0.1, 1, 10.0] |
|  |  | l1_ratio | [0.2, 0.4, 0.6, 0.8] |
|  |  | penalty | ['l1', 'l2', 'elasticnet'] |
|  |  | solver | ['lbfgs', 'newton-cg', 'saga'] |
|  | RF | n_estimators | [1000, 2000, 5000] |
|  |  | max_depth | [6, 8] |
|  |  | criterion | ['gini', 'entropy'] |
| Ensemble:  LR+RF+SVM | LR | max_iter | [100, 1000] |
|  |  | C | [0.0001, 0.001, 0.01, 0.1, 1, 10.0] |
|  |  | l1_ratio | [0.2, 0.4, 0.6, 0.8] |
|  |  | penalty | ['l1', 'l2', 'elasticnet'] |
|  |  | solver | ['lbfgs', 'newton-cg', 'saga'] |
|  | RF | n_estimators | [1000, 2000, 5000] |
|  |  | max_depth | [6, 8] |
|  |  | criterion | ['gini', 'entropy'] |
|  | SVM | kernel | ['linear', 'rbf'] |
|  |  | C | [0.001, 0.01, 0.1, 1] |
|  |  | gamma | [0.0001, 0.001, 0.01, 0.1] |
|  |  | degree | [2, 3, 4] |

**Supplementary Materials Table 3. List of non-optimised (i.e., fixed) hyperparameters for each machine learning model**

| **Classifier** | **Hyperparameter** | **Fixed value** |
| --- | --- | --- |
| LR | random_state | 38 |
| RF | random_state | 38 |
| SVM | probability | TRUE |
|  | random_state | 38 |
| Ensemble:LR+RF | voting | soft |
|  | random_state | 38 |
| Ensemble:LR+RF+SVM | Voting | soft |
|  | random_state | 38 |
|  | LR_algorithm | “auto” |
|  | SVM_probability | TRUE |

#### Features importance analysis with SHAP

We used the Python library shap v0.45.0 (SHAP) to analyses features importance during both training and test. SHAP is model-agnostic and hence can be applied on any machine learning models due to its versatile explainer functions. We used the KernelExplainer [32] to explain the LR, SVM, Ensemble:LR+RF, and Ensemble:LR+RF+SVM models. We used the TreeExplainer [33] to explain the RF model, which is designed for tree-based machine learning models. It is important to note that SHAP scores produced from different explainers are not on the same scale. Therefore, in this study, we only used SHAP scores to compare the importance of features within each model, not across models.

In addition to being model-agnostic, SHAP can be used at both local (i.e. sample) and global (i.e. dataset) levels. At the local level, SHAP produces a score for each feature of each sample representing its influence each patient to respond well or poorly towards ICI. At the global level, we can retrieve the absolute median SHAP score across all samples to explain the overall influence of each feature on a model. We leveraged this advantage of SHAP to study features importance in both training and test data.
